# Structural Basis of Non-Latent Signaling by the Anti-Müllerian Hormone Procomplex

**DOI:** 10.1101/2024.04.01.587627

**Authors:** James A Howard, Lucija Hok, Richard L Cate, Nathaniel J Sanford, Kaitlin N Hart, Edmund AE Leach, Alena S Bruening, David Pépin, Patricia K Donahoe, Thomas B Thompson

## Abstract

Most TGFβ family ligands exist as procomplexes consisting of a prodomain noncovalently bound to a growth factor (GF); Whereas some prodomains confer latency, the Anti-Müllerian Hormone (AMH) prodomain maintains a remarkably high affinity for the GF yet remains active. Using single particle EM methods, we show the AMH prodomain consists of two subdomains: a vestigial TGFβ prodomain-like fold and a novel, helical bundle GF-binding domain, the result of an exon insertion 450 million years ago, that engages both receptor epitopes. When associated with the prodomain, the AMH GF is distorted into a strained, open conformation whose closure upon bivalent binding of AMHR2 displaces the prodomain through a conformational shift mechanism to allow for signaling.

## Main Text

The Anti-Müllerian Hormone (AMH) is a glycoprotein signaling ligand with pivotal functions in reproductive biology. Through its mutually specific receptor, Anti-Müllerian Hormone Receptor Type II (AMHR2), AMH coordinates the regression of the Müllerian ducts during male fetal development and regulates follicle development in adult females^1,2^. As such, the dysregulation of this critical pathway is linked to well-characterized reproductive disorders in both sexes^3–6^. While AMH levels have been widely used as a measure of ovarian reserve for some time, recent studies have continued to investigate the potential of AMHR2 agonists as oncofertility or contraceptive agents^7–11^.

Like other TGFβ family members, AMH is synthesized as a large precursor consisting of an N-terminal prodomain and a C-terminal growth factor (GF)^12^. This paradigm is common among all subclasses of the family, including TGFβs, activins, GDFs, and BMPs. During synthesis, two AMH prodomain chains become disulfide-linked, similar to TGFβ1-3, and are subsequently cleaved from the GF, also a covalent dimer, by proprotein convertases^13,14^. The prodomains of TGFβ ligands are essential for proper folding and secretion of the GF, often persisting in a noncovalent procomplex with the GF following secretion^15,16^. For example, the GFs of TGFβ1 and GDF8 form a high-affinity prodomain-GF interaction that renders the GF latent until additional activation steps (i.e. mechanical force or proteolysis) liberate it for signaling^17–22^. Alternatively, for Activin A and BMP9, the procomplex interaction is weaker and does not interfere with signaling^23–25^. Regardless of these functional differences, the prodomains of most family members share general features and an overall similar fold often defined as a “TGFβ propeptide” domain. AMH, on the other hand, has evolved to have a substantially larger prodomain that aligns poorly with other family members. In addition, while the high affinity procomplex remains stable through secretion, it promotes biological activity rather than conferring latency^26–28^. Studies have shown that the procomplex form of AMH is required for the full regression of the Müllerian ducts as well as the inhibition of aromatase expression^26,29^. Despite its requirement for full bioactivity, the prodomain is broadly uncharacterized and the mechanism by which it maintains a high affinity for the GF without inhibiting cell signaling is unknown.

To elucidate the important molecular features of the AMH procomplex, we determined a cryo-EM structure of the binding interaction between the prodomain and the GF (**Fig. 1a**). Purified procomplex was combined with fab fragments of the 6E11 antibody to specifically target the prodomain-GF interface within AMH^30^. From one dataset, we generated two maps with and without imposed C2 symmetry, achieving global resolutions of 3.2 Å and 3.4 Å, respectively (**Extended Data Fig. 1 and Table 1**). While the C2 map contains a representation of the idealized full conformation for the procomplex, a symmetric dimer, continuous heterogeneity within the sample lead to a destructive pseudosymmetry which was mitigated by the C1 map. In both, the fab fragments and the core of the GF dimer are well ordered, whereas density and resolution both decline from ∼3 Å at the center of the procomplex to ∼5 Å at the outer prodomain helices due to innate flexibility (**Extended Data Fig. 2a,b**).

**Fig. 1.**
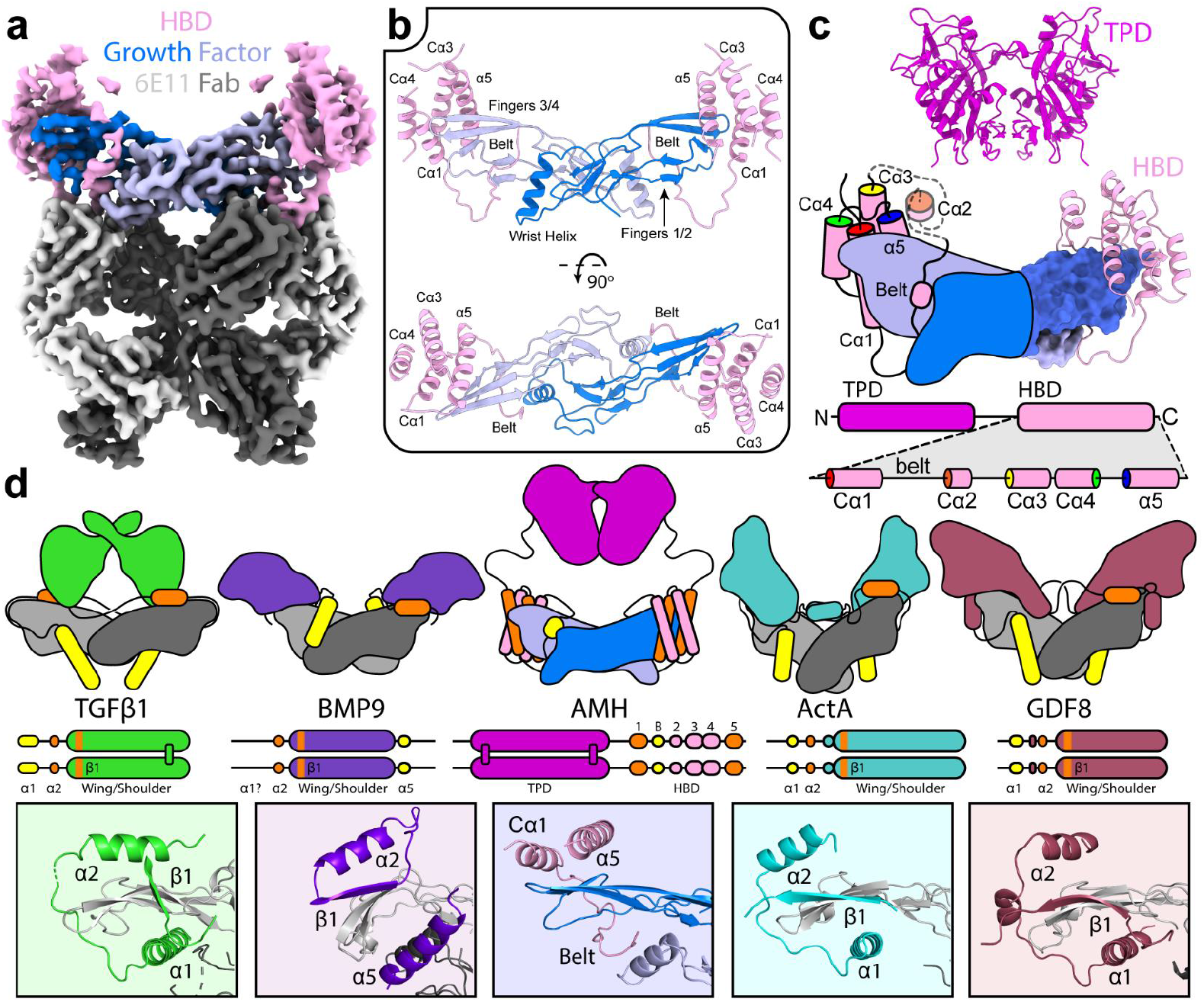
Structure of the AMH procomplex. **a**, Composite DeepEMhancer sharpened C1 map colored by protein components. b, Ribbon diagram shown for the full C2 dimer of the prodomain-GF structure, built by symmetry expansion of the C1 structure. c, Full AMH procomplex model generated using AlphaFold Multimer with structural annotation of the HBD. d, Cartoon, linear, and pro-GF fingertip interface comparison of structurally resolved prodomain subclasses. Cartoon representations are colored by proximity to type II (orange) and type I (yellow) binding sites. PDBs used are as follows: 5VQP (TGFβ1), 4YCG (BMP9), 5HLY (Activin A), and 5NTU (GDF8).

**Table 1.** Cryo-EM data collection, refinement, and validation statistics.

|  | C1 Map | C2 Map |
| --- | --- | --- |
| **Data collection and processing** |  |  |
| Magnification | 81,000 | 81,000 |
| Voltage (kV) | 300 | 300 |
| Electron exposure (e<sup>-</sup> Å<sup>-2</sup>) | 50.5 | 50.5 |
| Defocus range (μm) | −0.6 to −2.7 | −0.6 to −2.7 |
| Pixel size (Å) | 0.4495 | 0.4495 |
| Symmetry imposed | C1 | C2 |
| Initial particle images (no.) | 940,105 | 940,105 |
| Final particle images (no.) | 69,149 | 69,149 |
| Map resolution (Å) | 3.39 | 3.2 |
| FSC threshold | 0.143 | 0.143 |
| Map resolution range (Å) | 2.087 to 41.915 | 1.964 to 36.634 |
| **Refinement** |  |  |
| Initial model used (PDB code) | N/A | N/A |
| Model resolution (Å) | 3.4 | 3.2 |
| FSC threshold | 0.143 | 0.143 |
| Model resolution range (Å) | 3.1 to 3.9 | 2.8 to 3.7 |
| Map sharpening B factor (Å<sup>2</sup>) | −122.8 | −136.62 |
| **Model composition** |  |  |
| Nonhydrogen atoms | 8,877 | 8,876 |
| Protein residues | 1,169 | 1,165 |
| Ligands | N/A | N/A |
| **B factors (Å<sup>2</sup>)** |  |  |
| Protein | 124.7 | 132.4 |
| Ligand | N/A | N/A |
| **RMS<sup>a</sup> deviations** |  |  |
| Bond lengths (Å) | 0.004 | 0.003 |
| Bond angles (°) | 0.608 | 0.596 |
| **Validation** |  |  |
| MolProbity score | 1.57 | 1.59 |
| Clashscore | 4.15 | 4.15 |
| Poor rotamers (%) | 0.00 | 0.00 |
| **Ramachandran plot** |  |  |
| Favored (%) | 94.64 | 94.21 |
| Allowed (%) | 5.36 | 5.79 |
| Disallowed (%) | 0.00 | 0.00 |
| <sup>a</sup>RMS, root mean square deviation. N/A, not applicable. |  |  |

A near-complete atomic model was built for both the 6E11 fab and GF, while the prodomain was modeled more conservatively (**Extended Data Fig. 2c, Table 1**). We observed that the prodomain region resolved within our maps corresponded to a series of helices that are specific to the AMH prodomain and share no homology with any other TGFβ family member. This region, which we term the helical bundle domain (HBD), consists of a noncontiguous four-helix bundle which associates with the knuckles and fingertips of the GF (**Extended Data Fig. 2d,f**). A second interface is comprised of a “binding belt” that is formed from a loop that extends from the first helix (Cα1) of the bundle, runs through the inside of the GF fingers, and connects back to the bundle through Cα3 **(Fig. 1b,c and Extended Data Fig. 2e**). The helical bundle is completed by Cα4 and α5, the latter being named for its alignment to a more typical prodomain α5. Finally, the structure reveals that the 6E11 epitope occurs on the underside of the GF, stabilizing the procomplex form by closely contacting both the GF and prodomain.

Although the cryo-EM sample contained the full prodomain, only the HBD could be resolved, prompting us to utilize AlphaFold2 to generate a complete structural model. Our modeling results indicated the presence of a two-domain architecture within the AMH prodomain, which consists of an N-terminal TGFβ propeptide-like domain (TPD) connected by a proline-rich linker to the C-terminal HBD (**Fig. 1c and Extended Data Fig. 3a,b**). Furthermore, previous studies have shown that the AMH prodomain, which contains five cysteines, dimerizes via an intermolecular disulfide bond similar to the prodomains of TGFβ1-3^13,31^. Since our model indicates that the two prodomain chains dimerize via the TPD, we wanted to validate its accuracy by identifying the cysteines responsible for dimerization. Through mutational analysis, we identified that C55 and C241 form two intermolecular disulfide bonds linking two TPDs. Additionally, C103 and C188 form an intramolecular disulfide within the TPD, leaving C411 within the HBD unpaired (**Extended Data Fig. 3c–e**). Of note, disruption of the intermolecular disulfide bonds within the TPD did not significantly impact AMH secretion, unlike previously studied mutations or disruptions within this domain^32,33^.

The more general characteristics of the procomplex were investigated by small-angle X-ray scattering (SAXS) and molecular dynamics (MD) studies. From the SAXS data, a Kratky plot with two Gaussian peaks and a plateau at high q values that slowly decays to zero indicated that the AMH procomplex is a multidomain protein with interdomain flexibility (**Extended Data Fig. 4a,b**). From this data, we generated an ab initio bead model of the average procomplex in solution, into which we could easily dock the individual TPD dimer and the HBD-GF models (**Extended Data Fig. 4c**). From this starting position, the unstructured N-terminal extension and interdomain linker were rebuilt to generate a full, connected model of the AMH procomplex. MD simulation using this input showed that the interdomain linkers are the most flexible part of the prodomain and enable both axial and lateral movements of the TPD, resulting in an asymmetric, flattened structure of the procomplex (**Extended Data Fig. 4d**). The computationally derived radius of gyration for the equilibrated structure, Rg = 45.6 Å, closely agrees with the experimental value of 45.5 Å (**Extended Data Fig. 4e**). Further visualization by negative stain EM of prodomain complexes visually confirmed the presence of the predicted two-domain structure. Generated 2D classes of the AMH procomplex are consistent with an elongated HBD-GF complex connected to a dimerized TPD (**Extended Data Fig. 5a,b**). This two-domain structure was maintained within an unbound prodomain sample where, without the GF, the two HBDs closely associate (**Extended Data Fig. 5c,d**). When the 6E11 fab complex was visualized, the TPD dimer was only intermittently present in 2D classes, which supports that the flexibility of the tethered domain is what prevented its resolution within our cryo-EM structure (**Extended Data Fig. 5e,f**).

Using existing procomplex structures, we compared the binding mode of the AMH prodomain with experimentally derived profiles of TGFβ1, Activin A, GDF8, and BMP9 prodomains (**Fig. 1d**)^22,23,25,34^. In all cases, the prodomain component occludes both the concave type I receptor binding site, typically utilizing an α1 or α5 helix, as well as the fingertips and knuckle region of the convex type II receptor binding site, often using an α2 helix and shoulder domain. In lieu of an α1 or α5 helix, the type I site of the AMH GF is occupied by the more unstructured binding belt. The most extensive prodomain interface is at the knuckles of the type II site, where the AMH prodomain occupies a much broader surface using the Cα1 and α5 of the HBD to bind perpendicular to the β-sheet fingers, very distinct from the α2 helix of other classes which runs parallel to the GF fingers (**Fig. 1d and Extended Data Fig. 2d–f**).

Comparison to a modeled AMH-AMHR2-ALK2 ternary complex shows that the helical bundle engages the GF at the AMHR2 interface while the belt binds at the ALK2 epitope. The type I site within the procomplex is disrupted by the binding belt which occludes the nonpolar core of the concave surface, capping it at the bottom with several polar, charged residues. Meanwhile, using the same interface as AMHR2, the prodomain binds closer to fingertips while the receptor prefers a knuckle interaction (**Fig. 2a**)^35,36^. Interestingly, to allow for the shared use of the AMHR2 binding interface, the GF adopts an open conformation within the procomplex that differs from other GFs in the TGFβ superfamily, all of which present a more closed conformation similar to that observed in the AMHR2-bound structure (**Extended Data Fig. 6a,b)**. This is mainly visible in the 38° rotation of the fingertips when switching from the open and extended prodomain-bound form to the closed receptor-bound form (**Fig. 2b**). Examination of the structural features of this conformational shift reveals an inward movement of the GF fingers by ∼15 Å allowing them to form ionic interactions with the wrist helix of the opposite monomer, which could stabilize the closed conformation (**Extended Data Fig. 6c,d**). The transition from the open to the closed conformation occurs through a difference in the hydrogen bonding network within the GF β-sheets brought about by a 180° inversion of the V477 and L478 side chains on GF finger 2 (**Extended Data Fig. 6e,f**). An increase in the number of intramolecular contacts measured in MD simulations supports that the receptor-bound GF structure is more stable and suggests that the open conformation is likely more strained while bound by the HBD (**Extended Data Fig. 6f–g**).

**Fig. 2.**
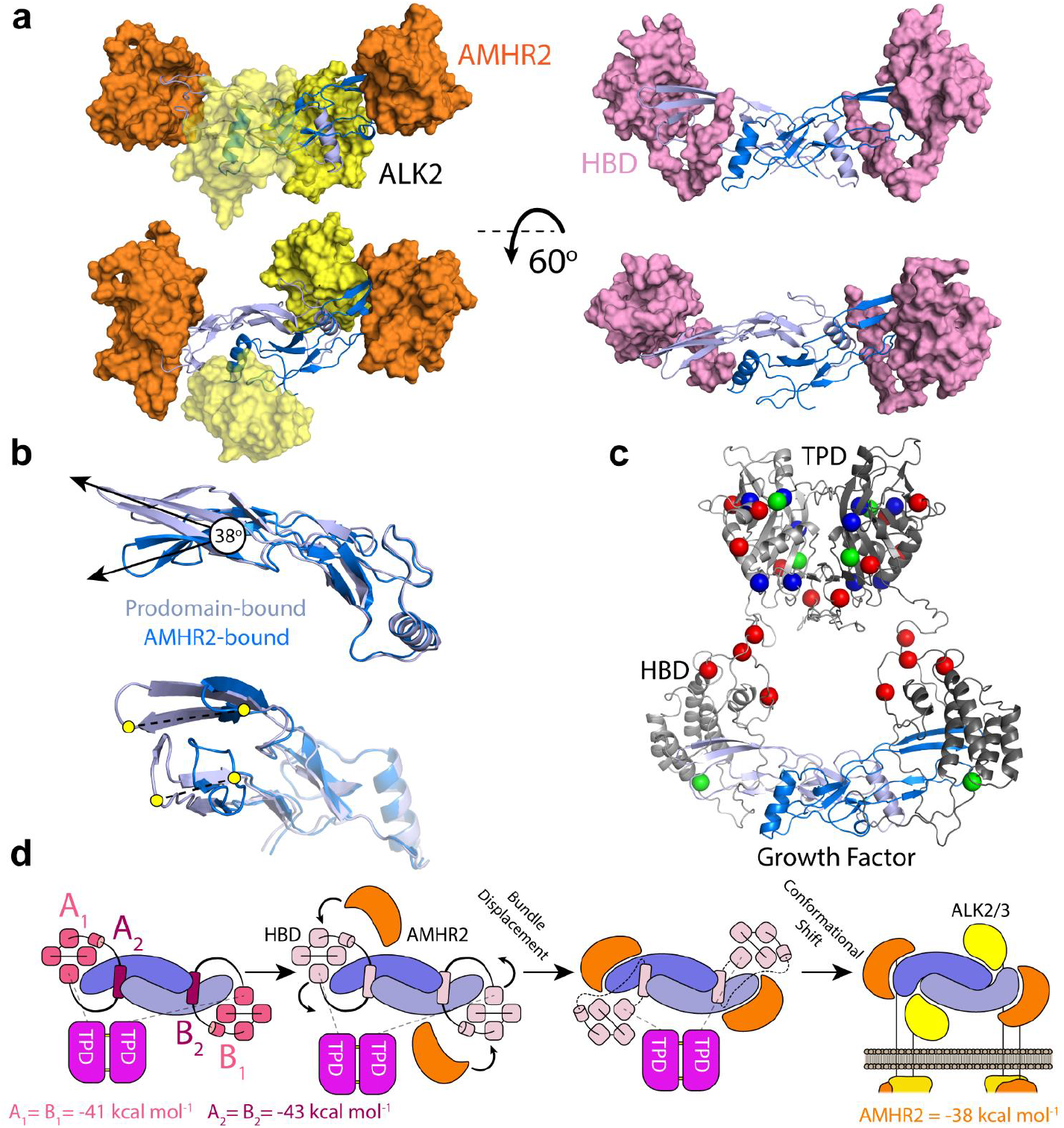
Functional consequences of the atypical prodomain architecture. **a**, Comparison of the modeled ternary complex of the AMH GF, AMHR2, and ALK2 with the cryo-EM structure of the AMH GF-HBD complex. **b**, Conformational shift within the GF from the open prodomain-bound state to the closed AMHR2-bound state. **c**, Annotation of residue positions with recurrent or characterized disease-associated mutations within the prodomain. PMDS are shown in blue, PCOS are shown in red, and PMDS/PCOS are shown in green. **d**, A tentative mechanism for displacement of the AMH prodomain involving displacement of the helical bundle and conformational shift within the GF to arrive at the active signaling complex. MM-GBSA binding enthalpies are shown in shades of pink on the left for helical bundle (A_1_, B_1_) and binding belt (A_2_, B_2_), as well as for AMHR2 on the right in orange (**Supplementary Table 1**).

As the HBD is an AMH-specific element among TGFβ prodomains, we were interested in its origins. We found that *amh* genes from all examined species, including the sex-linked *amhy* gene in certain teleost species, contain the HBD. Genetic alignments indicate that this domain arose before the last common ancestor to cartilaginous and bony fishes around 450 million years ago. This occurred as an exonic insertion within a primordial *amh* gene immediately upstream of the final exon encoding a prodomain α5 helix and the GF (**Extended Data Fig. 7a**). This separate HBD architecture is present within the *amh* gene of ray-finned fish, but in sharks, lobe-finned fishes, and tetrapods it has combined into one larger exon with the GF. Further modeling of 100 AMH homologs using AlphaFold2 showed consistency with the human structures (**Extended Data Fig. 7b–d**). It is interesting to note that the organization of secondary structure within the TPD matches the general architecture of a TGFβ family prodomain. It appears that this prodomain-like structure has been maintained with high conservation over the evolutionary history of AMH, especially the cysteine residues responsible for dimerization, despite losing GF binding function upon the divergence of the AMH gene (**Extended Data Fig. 7e**).

Using our cryo-EM structure, we could now map many pathogenic mutations which have previously been identified within the AMH prodomain (**Fig. 2c and Extended Data Fig. 8**). These focused on mutations associated with Persistent Müllerian Duct Syndrome (PMDS), a rare congenital difference of sexual development in the XY population, and Polycystic Ovary Syndrome (PCOS), a complex metabolic disease in the XX population characterized by various reproductive disruptions including aberrant AMH signaling^37,38^. Several PMDS and PCOS mutations including L339P, P352S, and P362S localize to the unresolved region of the HBD which has minimal if any interactions with the GF, suggesting a role for this region outside of direct GF binding. Others map to the binding interface of the HBD and most likely alter procomplex stability. For example, the prodomain mutation R302Q, which is found in both PMDS and PCOS, could disrupt a critical hydrogen bond between Cα1 of the HBD and L478 of GF finger 2. The A314G mutation, which occurs in certain cases of PMDS, is located in the binding belt and might disturb the backbone of the belt and affect the overall binding conformation. Also identified in patients with PMDS, the L426R mutation at the top of HBD α5 could interfere with the hydrophobic interactions formed on the interface with finger 3 and 4 of the GF and within the HBD. Most interestingly, it seems patient mutations are enriched within the TPD, which does not bind the GF. Previous data suggest that this domain may instead allosterically influence folding and dimerization of the GF^32,33^.

A unique feature of the AMH procomplex is that the prodomain maintains a high affinity for the GF, blocking its AMHR2 epitope, without rendering it latent. Consistent with previous studies, our procomplex structure supports that the prodomain must be fully displaced for AMH to signal, but not to bind its cognate receptor^27,28^. We see that the affinity of the prodomain is spread across four semi-independent interacting motifs consisting of two binding belts and two helical bundles which are covalently tethered through the TPD dimer. Through computational analysis, we found that each separate motif has an individual binding strength that is comparable to that of the receptor **(Fig. 2d and Extended Data Fig. 9a)**. Moreso, dynamics of the prodomain monitored over time indicate a more fluid helical bundle, compared to a more stable belt, which could promote its partial displacement by the receptor (**Extended Data Fig. 9b–d**). Thus, an encounter with one AMHR2 on the cell surface will only displace the helical bundle on one side, allowing for the prodomain to remain bound to the GF on that same side through the binding belt. Engagement of a second AMHR2 on the cell surface then leads to a conformational change in the GF, inducing a closing motion within the fingers which facilitates full prodomain displacement. This rearrangement frees the GF from the belt and allows the type I receptor to bind the closed GF, which we believe is represented within the crystal structure, to form the signaling complex. It is possible that the directionality of this transition is supported by the stabilization of the GF itself, which moves from a more open and strained conformation within the procomplex to a stable, closed conformation when bound by AMHR2 (**Extended Data Fig. 6g**). This mechanism is consistent with the concentration dependence of prodomain displacement observed when procomplex binds to AMHR2 on a surface; displacement is only observed at low AMH concentrations where bivalent binding to AMHR2 is favored^28^. Nevertheless, the full mechanism underlying prodomain displacement, and what drives the conformational shift, requires further investigation.

As these studies show, AMH is an atypical molecule which exists outside of the paradigms defined for other TGFβ family members. Its evolutionarily divergent two-domain structure makes this prodomain incomparable to even its nearest homologs. Herein, we have elucidated an entirely new binding mode for the AMH prodomain which begins to explain the functional paradox of how it can maintain high affinity for the GF without conferring latency. These findings are especially important when considering the functional necessity of the mutant-prone prodomain in vivo and understanding the mechanism by which it promotes signaling. Owing to the critical importance of the prodomain for bioactivity, our structural model provides a framework for further investigation of the AMH signaling pathway, ultimately striving for the development of protein therapeutics with enhanced pharmacological properties.

## Methods

### Protein purification and complex generation

#### AMH Procomplex

Recombinant AMH protein was expressed and purified as previously reported^10,35,36,39^. Enhanced AMH protein containing an albumin leader sequence and Q450R mutation for enhanced cleavage (LR-MIS) was produced using a stably transfected Chinese hamster ovary cell line. Conditioned media was initially purified by affinity chromatography using an N-hydroxysuccinimide (NHS) column (Cytiva) primary amine coupled with full-length 6E11 antibody. A second size-exclusion chromatography step using a Superdex Increase 200 column (Cytiva) was used to separate dimeric AMH from oligomer species.

#### mAb 6E11

Anti-human MIS monoclonal antibody (clone 6E11)^30^ was produced by stably transfected B-cell hybridoma cell line. Conditioned media was then purified by affinity chromatography using HiTrap protein A HP column (Cytiva).

#### AMH Prodomain

The LR-MIS expression construct was modified using site-directed mutagenesis to add a stop codon following R451 within the human AMH protein. Recombinant prodomain protein was produced by transient transfection of Expi293 cells at a concentration of 1 μg mL^−1^ cell culture at a cell count of 2.5×10^6^ cells mL^−1^. Conditioned media was collected after 3 days shaking at 37 °C and loaded onto a HiTrap SP HP cation exchange chromatography column (Cytiva), washed, and eluted using a salt gradient. A second size-exclusion chromatography step using a Superdex Increase 200 column (Cytiva) was used to separate dimeric prodomain from oligomer species and other contaminants.

#### AMH-6E11 complex generation

Full-length 6E11 antibody was digested over 4 hours at 37 °C using immobilized ficin agarose resin (Thermo Fisher) and then clarified by size-exclusion chromatography using a Superdex Increase 200 column (Cytiva) to isolate fab fragments. Excess 6E11 fab was then combined with dimeric AMH procomplex in sizing buffer (10 mM MES, 150 mM NaCl, pH = 5) and incubated for 1 hour while rocking at 4 °C. The resulting complex was then concentrated and isolated using a Superdex Increase 200 column (Cytiva) before immediate use in structural applications.

### Cryo-EM Studies

#### Cryo-EM Sample Preparation and Data Collection

AMH-6E11 complex was prepared for cryo-EM studies by concentration to 0.6 mg mL^−1^ in 10 mM MES, 150 mM NaCl, pH = 5.5. Holey copper Quantifoil grids (Ted Pella) with 300 mesh and R1.2/1.3 hole spacing were glow discharged for 30 s at 15 mA and 0.4 mbar using the Pelco EasiGlow. 4 μL of sample was applied and the grid was blotted for 5 seconds at 4 °C and 95% relative humidity then plunge-frozen with liquid ethane using a Vitrobot (ThermoFisher). Samples were clipped and screened by Glacios transmission electron microscope before 5,276 movies were collected on a Titan Krios transmission electron microscope at the Center for Electron Microscopy and Analysis (CEMAS) at Ohio State University. The movies were imaged in counted super-resolution with a nominal resolution of 0.4495 Å per pixel using a Gatan K3 detector with energy filter and Cs corrector. Each movie had a total electron dose of 50.5 e Å^−2^ and so was subdivided into 50 frames.

#### Cryo-EM Data Processing

Data Processing was done using Cryosparc V4.4. Briefly, Movies were motion and CTF corrected using Cryosparc’s patch implementations. 500 Micrographs were used to generate an initial structure from which 2D templates were made. Particles were then picked from all 5,276 micrographs using these templates, resulting in 4,105,575 particle picks. Poor quality micrographs were removed and a total of 3,486,824 particles were extracted from 5,000 micrographs at a box size of 600 pixels, which was then Fourier cropped to 200 pixels for efficient processing. Multiple rounds of 2D classification and ab initio model generation were performed to generate 10 classes for a heterogeneous refinement with imposed C2 symmetry (one “real” class and 9 “junk” classes). Particles from the “real” class representing ∼27% of the total extracted particles (940,105) were expanded about a C2 symmetry axis (1,880,210) and used for a local refinement. The refined map and particles were then subjected to 3D variability analysis with a focus mask on the prodomain portion and divided into 10 variability clusters. Cluster 8, which comprised only 5.72% of the particles, contained the most prodomain density and thus was selected for further refinement. Particles from cluster 8 underwent reference-based motion correction and re-extraction at full resolution (box size = 600 px). After duplicate particles were removed, a global homogenous refinement was performed (94,363) which was then subdivided into 4 3D classes, each representing ∼25% of the total particle count. Class 1, which had the lowest global resolution, was excluded and the remaining 69,149 particles were used in a final global homogenous refinement to generate our C1 map with a global, masked resolution of 3.39 Å. The C2 map was generated using an additional homogenous refinement with imposed C2 symmetry and achieved a global, masked resolution of 3.20 Å.

#### Model building, Refinement, and Validation

Initial identification of map components was done using ModelAngelo with supplied FASTA sequences of the full sample protein (Full-length AMH and 6E11 fab). Once the prodomain region was confirmed to be the HBD, AF2 models of the GF-HBD complex and the 6E11 fab were docked into the map density. These models were then further trimmed and relaxed into the respective maps using ISOLDE and ChimeraX. Iterative rounds of real space refinement within Phenix followed by manual adjustment within Coot or using ISOLDE and ChimeraX resulted in a near-complete model of the 6E11 fab and AMH GF, with a conservative model of the HBD within which loops were excluded or side chains were truncated in regions without sufficient density or resolvability. Validations were derived from Phenix’s validation package as well as the wwPDB validation server. We have done our best to incorporate and conform to the recommendations and guidelines established by other experts^40^.

### Molecular Modeling

Models of human AMH were generated with ColabFold v1.5.5 (AlphaFold2 using MMseqs2)^41^. The full HBD:GF model was made using the cryo-EM structure as a template, while the TPD dimer was made without templates. Models for alternative species *amh* genes (including *amhy*) were generated on a local install of AlphaFold 2.3 as separate TPD dimer and HBD-GF structures^42^.

### Cysteine Mutagenesis

Full-length AMH (LR-MIS) was modified by site-directed mutagenesis to generate an initial 5 single cysteine-to-serine mutant constructs which targeted each cysteine residue within the prodomain. The signaling activity of these constructs was tested in the previously described AMH-responsive BRE-luciferase reporter assay system^35^. 50 ng of expression plasmid containing each construct was transiently transfected along with ALK2, AMHR2, and BRE-Luc, and the fold change in luminescence was analyzed following an 18-hour incubation. Mutations which maintained signaling activity (C55S, C241S, C411S) were combined by further site-directed mutagenesis to generate 4 more combinations. Eight 30 mL Expi293 cell cultures were transiently transfected with expression constructs containing wild-type or mutant AMH which were harvested after 72 hours. Conditioned media was loaded onto 4-20% SDS gels under non-reducing or reducing conditions, then analyzed by western blot using a polyclonal detection antibody directed against the prodomain (OAAB05640, Aviva Systems Biology).

### Small-angle X-ray Scattering

SAXS data of AMH were collected using SIBYLS mail-in SAXS service as previously described^43–46^. In brief, LR-MIS protein was prepared in PBS at 2 mg mL^−1^ and mailed to the SIBYLS beamline at 4 °C. The sample was isolated over a Shodex 803 SEC column to ensure a disaggregated sample immediately followed by SAXS measurements. Data were analyzed using PRIMUS to generate log and Kratky plots, distance distribution, and radius of gyration. DAMMIF was used within PRIMUS to generate an average P1 bead model of the procomplex.

### Negative Stain Electron Microscopy

400 mesh Cu formvar carbon grids (Ted Pella) were glow discharged for 30 s at 15 mA at 0.4 mbar using the Pelco EasiGlow. Procomplex, prodomain, and 6E11-procomplex samples were diluted to 5 μg mL^−1^ in 10 mM MES, 150 mM NaCl, pH = 5 before staining with 0.75% Uranium formate. 150, 161, and 75 micrographs were collected each for the procomplex, prodomain, and 6E11-procomplex samples, then processed using Relion 4 into 2D classes.

### AMH Evolution

The AMH gene structure in a particular species was determined by searching the RefSeq Genome database for that species with the AMH protein sequence for that species using tblastn (National Center for Biotechnology Information).

### Computational Analyses

#### Molecular Dynamics Simulations (MDS)

##### 1.1. AMH procomplex

Structures of the AMH bound to the prodomain containing both TPD and HBD domains by linkers were generated using AlphaFold2 (AF2) components and connected using ISOLDE within ChimeraX^42,47,48^. Three structurally distinct models – *i*) model01 with TPD placed above the HBD-GF complex containing less structured linkers; *ii*) model02 with the same spatial relation of domains as the previous one, but with more structured linkers; and *iii*) model03 with TPD located below the HBD-GF complex – were chosen as starting structures for MD simulations. Protonation states of ionizable amino acid residues were estimated by PROPKA 3.1 and by visual inspection of their hydrogen-bonding patterns, which prompted us to account for all amino acid residues in their typical protonation states^49^. Models were solvated in 5-Å-octahedral boxes and neutralized by 6 Cl^−^ anions, and subsequently parameterized using the AMBER ff14SB force field for proteins and the TIP3P model for water^50,51^. These were submitted to geometry optimization in AMBER16 with periodic boundary conditions in all directions^51^. The optimized systems were gradually heated from 0 to 300 K and equilibrated during 30 ps using *NVT* conditions, followed by productive and unconstrained MD simulations of 400 ns by employing a time step of 2 fs. The pressure of 1 atm and temperature of 300 K were kept constant using Berendsen barostat and Langevin thermostat, respectively, with a 1 ps^−1^ collision frequency^52,53^. The long-range electrostatic interactions were calculated employing the Particle Mesh Ewald method and were updated in every second step, while the non-bonded interactions were truncated at 11 Å^54^.

CPPTRAJ program, implemented in AMBER, was used for processing coordinate trajectories and data files^55^. Root Mean Square Deviation (RMSD) analysis was performed on backbone atoms of the entire procomplex, as well as individual parts of the prodomains: TPD, linkers, and HBD. 3D histograms of the procomplex position during 400 ns-MDS were generated using the *grid* command of CPPTRAJ and shown along with the average structures for each model. The theoretical SAXS curves were computed utilizing the saxs_md program within AMBER, according to the theory of excess electron density^56^. Water molecules within 5 Å of the protein were explicitly included in the calculation, which is consistent with previously modeled multidomain proteins^57^. To accurately estimate the electron density of the bulk solvent, we ran a 30-ns-long MD simulation involving 59,109 water molecules at the *NVT* ensemble in a periodic octahedral box with dimensions matching solute systems.

##### 1.2. Other Complexes

Initial structure for the procomplex involving only HBD was obtained by AF2 modeling based on our determined cryo-EM structure. The crystal structure of AMH bound to AMHR2(ECD) deposited under PDB ID: 7L0J was used as a starting point for MDS^36^. For both systems, firstly, the protonation states were determined by PROPKA 3.1^49^, after which complexes were solvated in a truncated octahedral box of TIP3P water molecules spanning 8 Å-thick buffer and neutralized by corresponding number of counterions: 10 Cl^−^ for the procomplex, while the AMH-AMHR2 complex already had a net charge of 0. Systems were parameterized using the AMBER ff14SB force field, and submitted to molecular dynamics simulations of 300 ns using the same setup as previously described^51^.

All simulations were run in five replicas, and the analysis of the procomplex and AMH-AMHR2 simulations was performed separately for each monomer in each of the five replicates, while the results were reported as average values. For the Root Mean Square Deviation (RMSD) analysis only backbone atoms were included, while Root Mean Square Fluctuation (RMSF) analysis was performed on side chain atoms as well. For the hydrogen bond calculations, the following criteria were used: donor-H···acceptor ∠ = 135°−180°, donor···acceptor distance = < 3 Å. The corresponding binding enthalpies were calculated on 3000 frames from the last 30 ns of simulations using the MM-GBSA protocol and then decomposed into specific residue contributions on a *per-residue* basis according to the previously established procedure^32,58,59^.

#### Umbrella Sampling Simulations

Umbrella sampling simulations were performed on the following systems: *i*) AMH with one HBD bound at the type II site; *ii*) AMH with one HBD without binding belt [amino acid residues: 303−331] bound at the type II site; and *iii*) AMH with one AMHR2(ECD) bound at the type II site, utilizing GROMACS 2022.4^60^ The simulations were set up in 9.0 nm x 22.0 nm x 9.0 nm, 9.0 nm x 20.0 nm x 9.0 nm, and 16.0 nm x 8.0 nm x 8.0 nm rectangular boxes, respectively, employing periodic boundary conditions in all directions. The systems were oriented so that the prodomains were aligned with the y-axis, and the receptor with the x-axis, along which they were pulled out during simulations. All proteins were solvated with TIP3P water molecules, while 7 Cl^−^, 7 Cl^−^, and 2 Cl^−^ ions for the procomplex, the procomplex without belt and the AMH-AMHR2 complex, respectively, were added to keep the overall charge of the simulation boxes neutral. The equations of motion were integrated with time steps of 2 fs. The buffered Verlet neighbor list was used to evaluate the Lennard-Jones (6,12) interactions up to a distance cutoff of 1.4 nm, while long-range electrostatic interactions were treated by the Particle Mesh Ewald method using a Fourier spacing of 0.12 nm^54^. The temperature was kept at 298.15 K using the Nosé-Hoover thermostat with a 1.0 ps^−1^ coupling time constant^61^. For constant 1 bar pressure, isotropic Parrinello-Rahman barostat was employed with 2.0 ps^−1^ coupling time constant and compressibility for water of 4.5 · 10^−5^/bar^62^. The prodomains and the receptor were pulled by a harmonic spring with force constant of 2,000 kJ mol^-1^ nm^-2^, moving away from the center of mass of the GF at constant velocity of 0.01 nm ps^-1^. To prevent the GF from being pulled together with the prodomains or receptor, a harmonic position restraint with a force constant 3,000 kJ mol^-1^ nm^-2^ was imposed on amino acid residues 505– 526 on the monomer not in direct contact with the prodomain or receptor. For subsequent umbrella sampling simulations, snapshots were extracted from the pulling simulations such that y-component of the GF-HBD distance or x-component of the GF-AMHR2 distance increases by 0.1 nm between neighboring windows, accounting for a total ∼60 windows. For each window, 50 ns of MD sampling was performed using the same parameters as described earlier (force constant of 2,000 kJ mol^-1^ nm^-2^, velocity of 0.00 nm ps^-1^). In case it was necessary, additional windows with the force constant of 5,000 kJ mol^-1^ nm^-2^ were added to ensure the necessary overlap of the histograms (**Supplementary Fig. 1**). The curves of the potential of mean force (PMF) were calculated from the last 40 ns of windows simulations by the Weighted histogram analysis method (WHAM), using the *gmx wham* GROMACS tool^63^. The statistical error of the PMFs was estimated via Bayesian bootstrap analysis with 100 bootstraps and plotted together with the average profile.

### Visualization

Figures and alignments were generated using ChimeraX and the Pymol package. The map in figure 1 was sharpened using DeepEMhancer’s high resolution model^64^.

## Reporting summary

Further information on results is available in the Supplementary Information.

## Acknowledgements

This work was funded by an R01 from the Eunice Kennedy Shriver National Institute of Child Health and Human Development (NICHD), R01HD105818. JAH is supported by the National Institute of Environmental Health Sciences, T32ES07250. LH thanks the University Computing Centre (SRCE) at the University of Zagreb, Croatia and the Advanced Research Computing (ARC) center at the University of Cincinnati, Cincinnati, OH, USA for providing computing resources. DP is supported by an R01 from the NICHD, 1R01HD102014. We thank Desirée Benefield at the Center for Advanced Structural Biology (CASB) at the University of Cincinnati and Yoshie Narui at the Center for Electron Microscopy and Analysis (CEMAS) at Ohio State University for their single-particle EM support. JAH thanks the Cold Spring Harbor Course on Cryo-Electron Microscopy, as well as Jennifer Cash, Gabriel Lander, and Justin Kollman, for their training and advice on single particle EM techniques.

## Author contributions

JAH, KNH, TBT conceived and designed the research. JAH, NJS, KNH, EAEL, and ASB performed experimental work. LH performed simulations and computational analyses. RLC performed evolutionary analyses. JAH, PKD, DP, RLC, and TBT interpreted data. JAH wrote the manuscript. All authors were involved in revising the manuscript.

---

**Extended Data Fig. 1.**
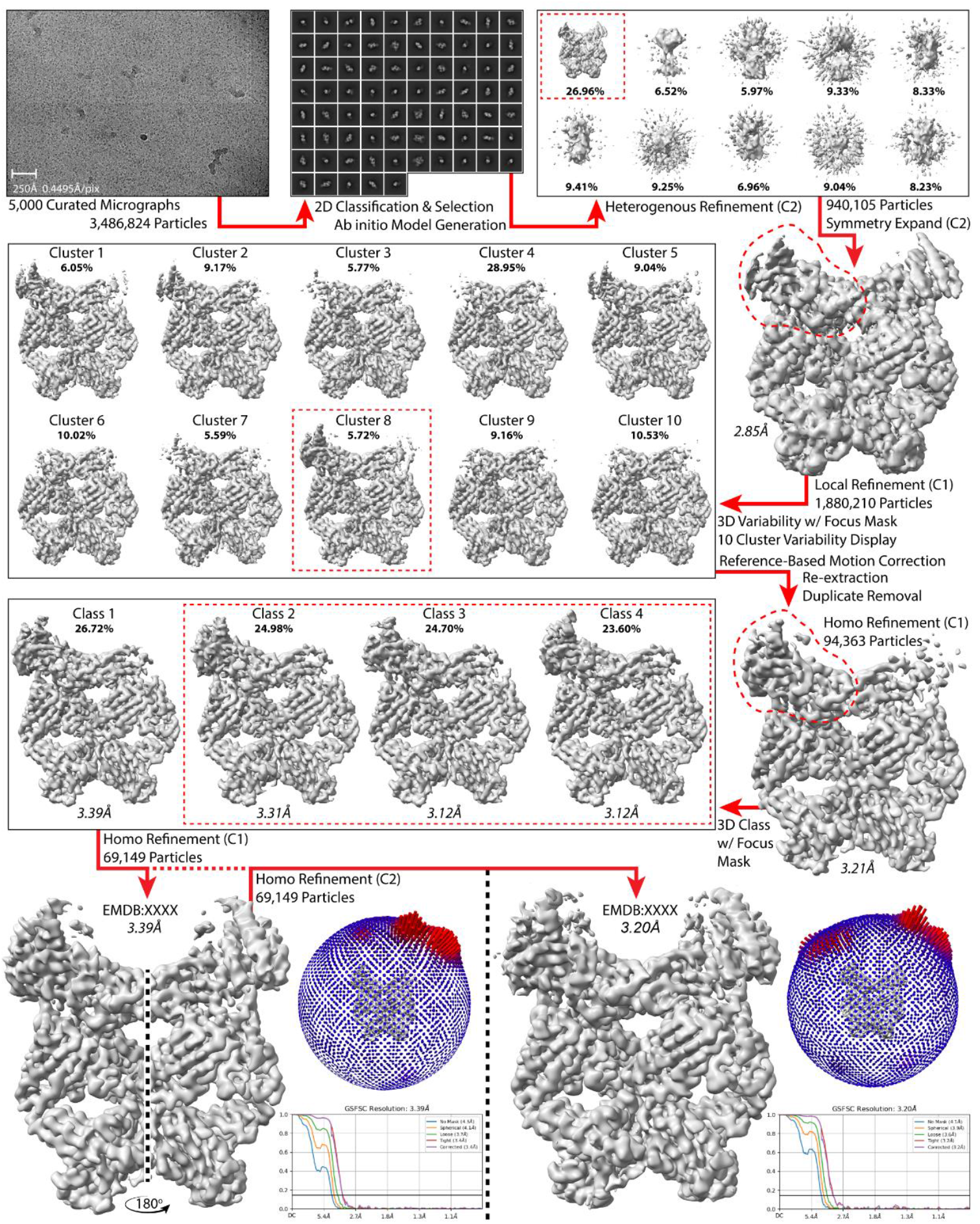
Cryo-EM processing workflow. Simplified scheme of data processing using CryoSparc. Red dashes indicate focus masks and selected clusters/classes. C1 map is shown with the resolved prodomain reflected about the central axis.

**Extended Data Fig. 2.**
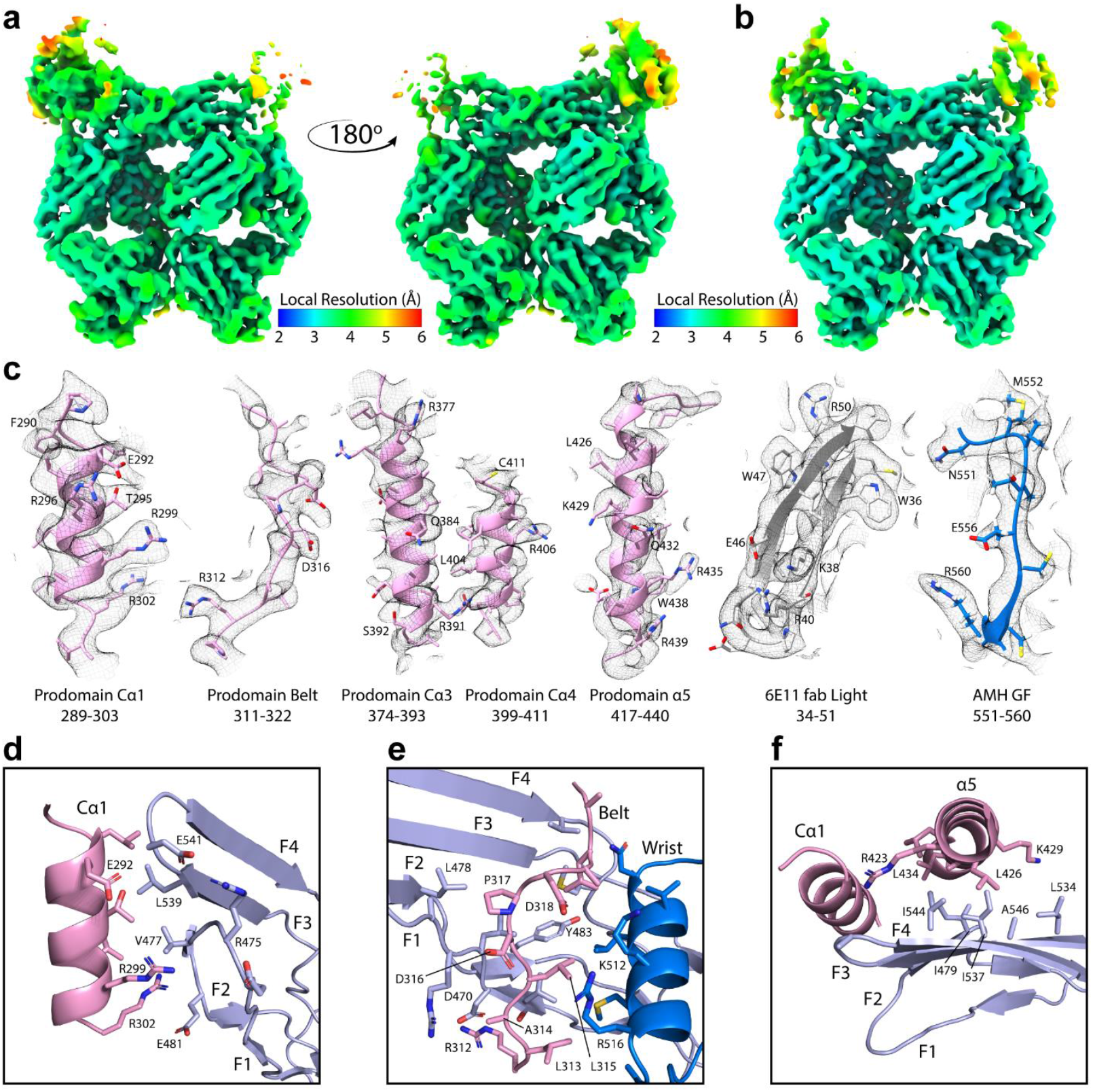
Cryo-EM validation and example density. **a, b**, Local resolution of C1 and C2 maps, respectively. **c**, Representative cryo-EM density fits for prodomain, fab, and GF within the *B* factor sharpened C1 map at a contour of 0.025. **d−f**, Molecular interactions at the prodomain-GF interface between: Cα1 helix and fingertips, binding belt and concave region, and Cα1/α5 helices and convex region.

**Extended Data Fig. 3.**
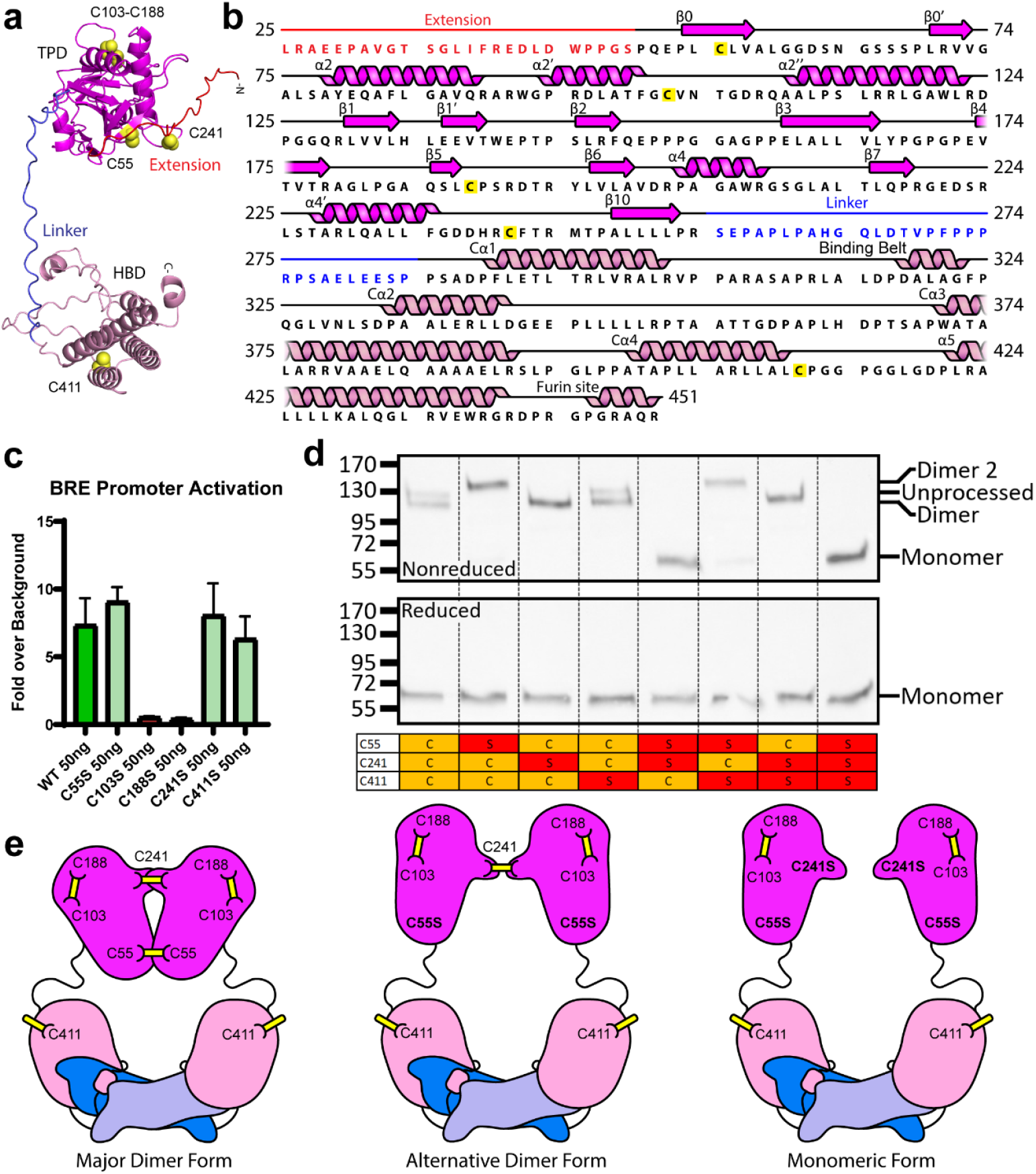
Molecular modeling and mutagenesis. **a**, AlphaFold monomer prediction of the AMH prodomain with cysteines shown. **b**, Prodomain sequence annotated with predicted secondary structure. TPD is labeled relative to established features of TGFβ1. **c**, BRE luciferase assay results using transfected AMH with single cysteine to serine point mutations within the prodomain. **d**, Reduced and nonreduced western blots probed for prodomain showing differences in structure and dimerization as a result of combinatorial cysteine to serine point mutations. **e**, Predicted architecture of different AMH forms.

**Extended Data Fig. 4.**
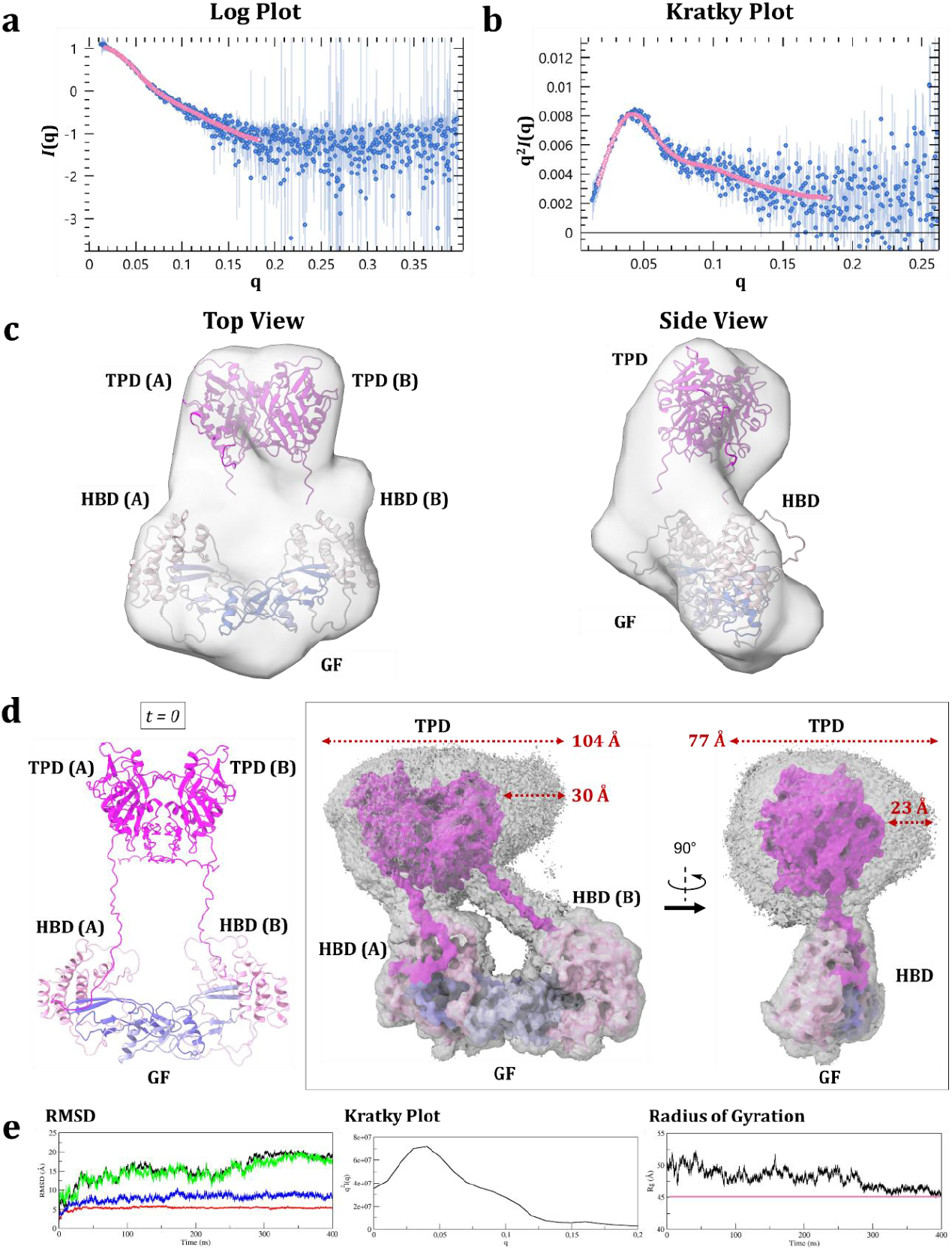
Small angle X-ray scattering analysis and global orientation of AMH procomplex. **a, b**, Kratky and log plots of AMH procomplex with fit line of distance distribution analysis shown. **c**, DAMMIF ab initio bead model generated from SAXS data with docked AMH components. **d**, Starting model from AlphaFold2-generated components for MD simulation with unstructured regions rebuilt (left). 3D histograms of the procomplex position during the MD simulation shown together with the average structure as a reference (right). HBDs are colored in pink and TPDs in magenta, while monomers in the GF are represented in different shades of blue. **e**, RMSD graph shown with a black line that signifies the backbone atoms of the entire procomplex (left). Red, green, and blue lines pertain to individual parts of the prodomain: TPD, linkers, and HBD, respectively. Kratky plot from the SAXS_MD analysis (middle). The radius of gyration during the MD simulation with the evolving value over 400 ns shown in black and the experimental value of 45.5 Å shown as a pink line (right). Results from MD simulations of two additional, structurally distinct models are given in **Supplementary Fig. 2**.

**Extended Data Fig. 5.**
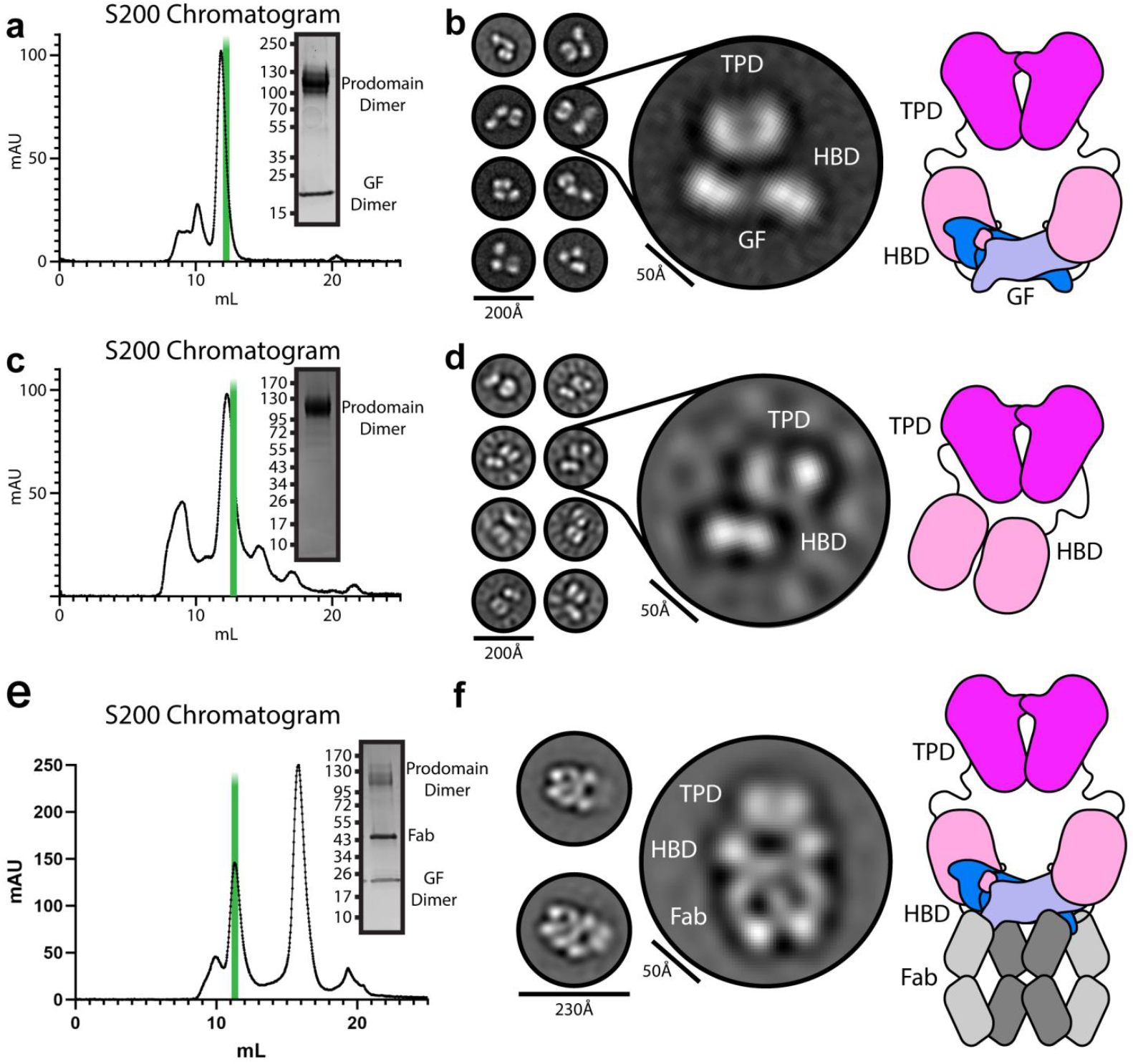
Negative stain EM of AMH complexes. **a,b**, SEC chromatogram with SDS gel and 2D classes of the AMH procomplex with cartoon model. **c,d**, SEC chromatogram with SDS gel and 2D classes of the AMH prodomain with cartoon model. **e,f**, SEC chromatogram with SDS gel and 2D classes of the AMH procomplex bound by 6E11 fab with cartoon model. Green bars in **a, b**, and **c** represent the fraction shown in the SDS gel inset and used in downstream structural applications.

**Extended Data Fig. 6.**
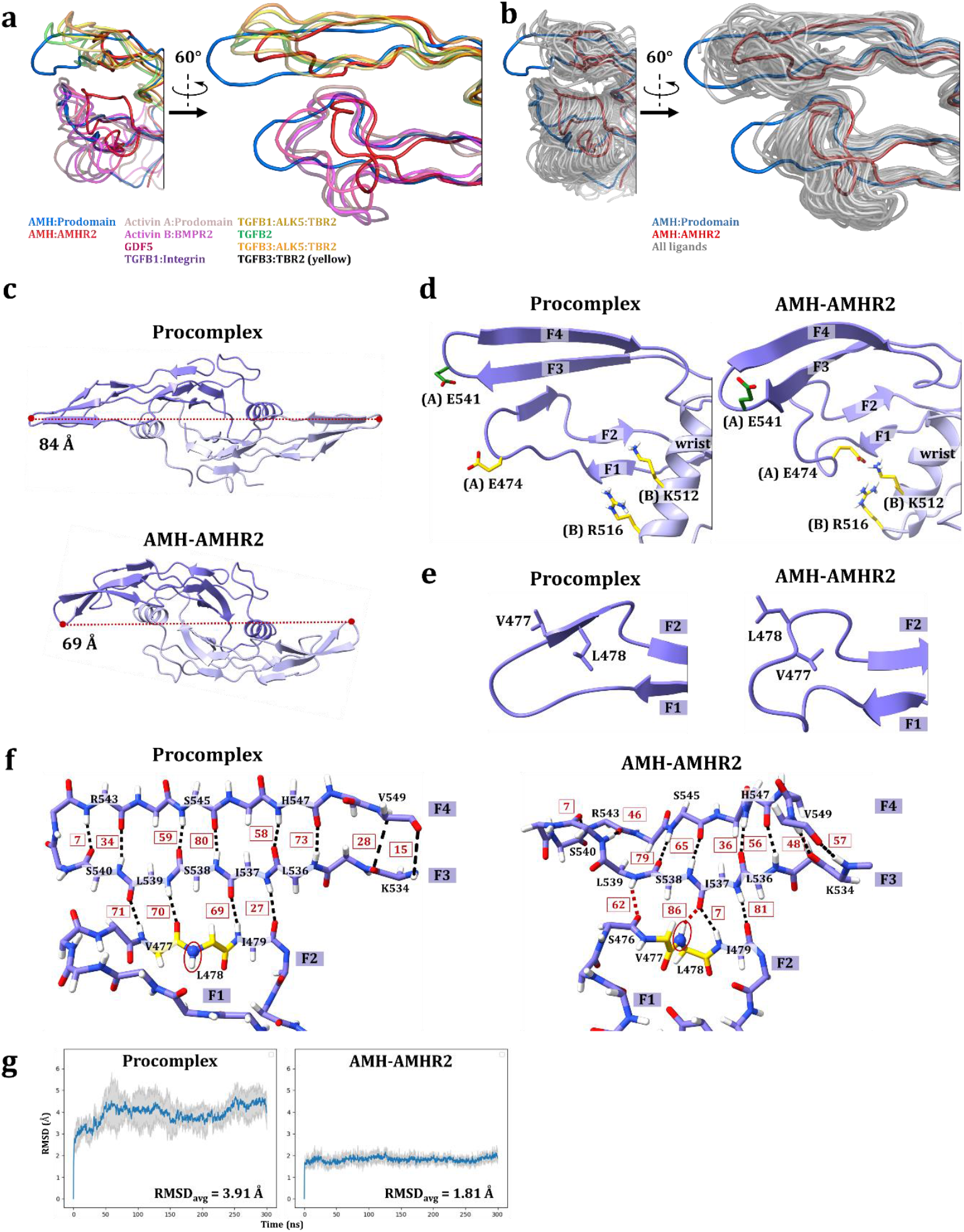
Comparison of GF conformations in open prodomain-bound form and closed receptor-bound form. **a**, Comparison of AMH GF structures in the procomplex and the AMHR2-bound form to the most similar conformations for fingers 1/2 and fingers 3/4 by backbone RMSD. Colors of the transparent ribbon representation correspond to the respective label for that structure. **b**, Overlay of the structures of all GFs from the TGFβ superfamily that are deposited in the PDB database with the AMH GF from the procomplex and the AMHR2-bound form. **c**, On the top, the open conformation of AMH in the procomplex as seen from the 84 Å distance between the E542 Cα-atoms in the opposite monomers, compared to the closed conformation in the AMH-AMHR2 complex on the bottom with a distance of 69 Å. **d**, On the left, downward orientation of the E541 side chain on fingertips 3/4 and the absence of ionic interactions between (A)E474 on finger 1 and (B)K512/(B)R516 on the wrist of the opposite GF monomer bound to the HBD. On the right, upward orientation of the E541 and the presence of the aforementioned interactions in the GF bound to the AMHR2. **e**, Opposite V477-L478 orientation observed in the GF bound to the HBD (on the left) or to the AMHR2 (on the right). **f**, On the left, linear hydrogen bonds between fingers 2/3 and 3/4 of GF forming three antiparallel β-sheets in the procomplex form. On the right, loss of the third β-sheet in finger 2 caused by the V477-L478 inversion in the GF bound to the AMHR2. The frequency of each hydrogen bond formation in procomplex and AMHR2-complex during MDS is given as a percentage calculated from the number of 150,000 frames. The loss of L539···V477 contacts is compensated by newly-created bonds between L539···S476 and I537···L478, as the measured number of hydrogen bonds formed between fingers 3 and 4 increased by 10% (**Supplementary Tables 2** and **3**). **g**, The RMSD graph of the GF backbone atoms on the left shows higher flexibility in the procomplex compared to the rigid and conformationally stable AMH-AMHR2 complex on the right (**Supplementary Table 4**).

**Extended Data Fig. 7.**
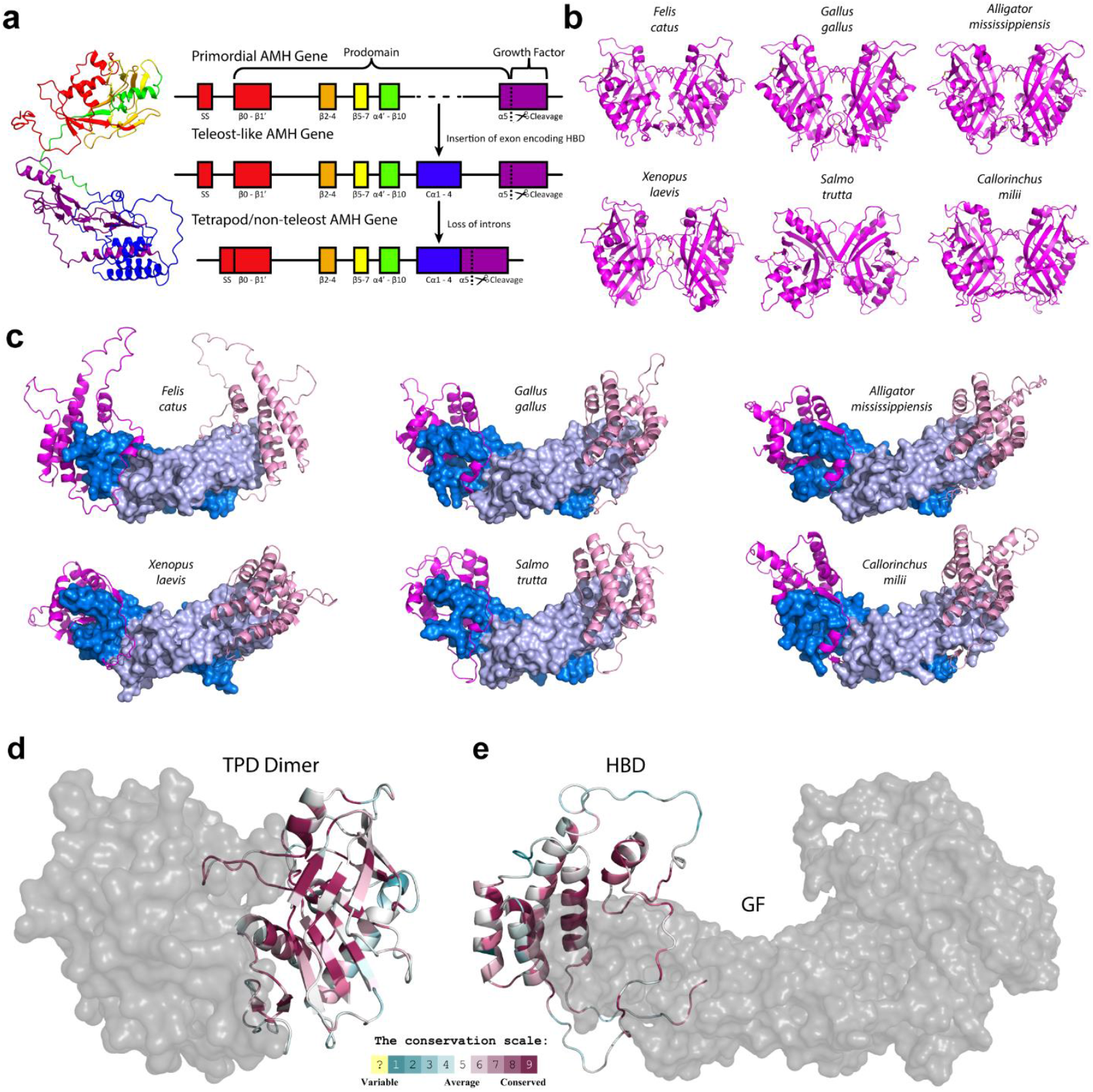
Evolution of AMH Prodomain. **a**, AlphaFold monomer model of AMH and gene annotation for AMH within tetrapods, teleost fish, and the hypothetical primordial TGFβ-like gene. **b**, TPD dimer models generated using AlphaFold Multimer for six representative species. **c**, HBD-GF models generated using AlphaFold2 Multimer for six representative species. **d, e**, ConSurf evolutionary conservation profiles of AMH TPD and HBD.

**Extended Data Fig. 8.**
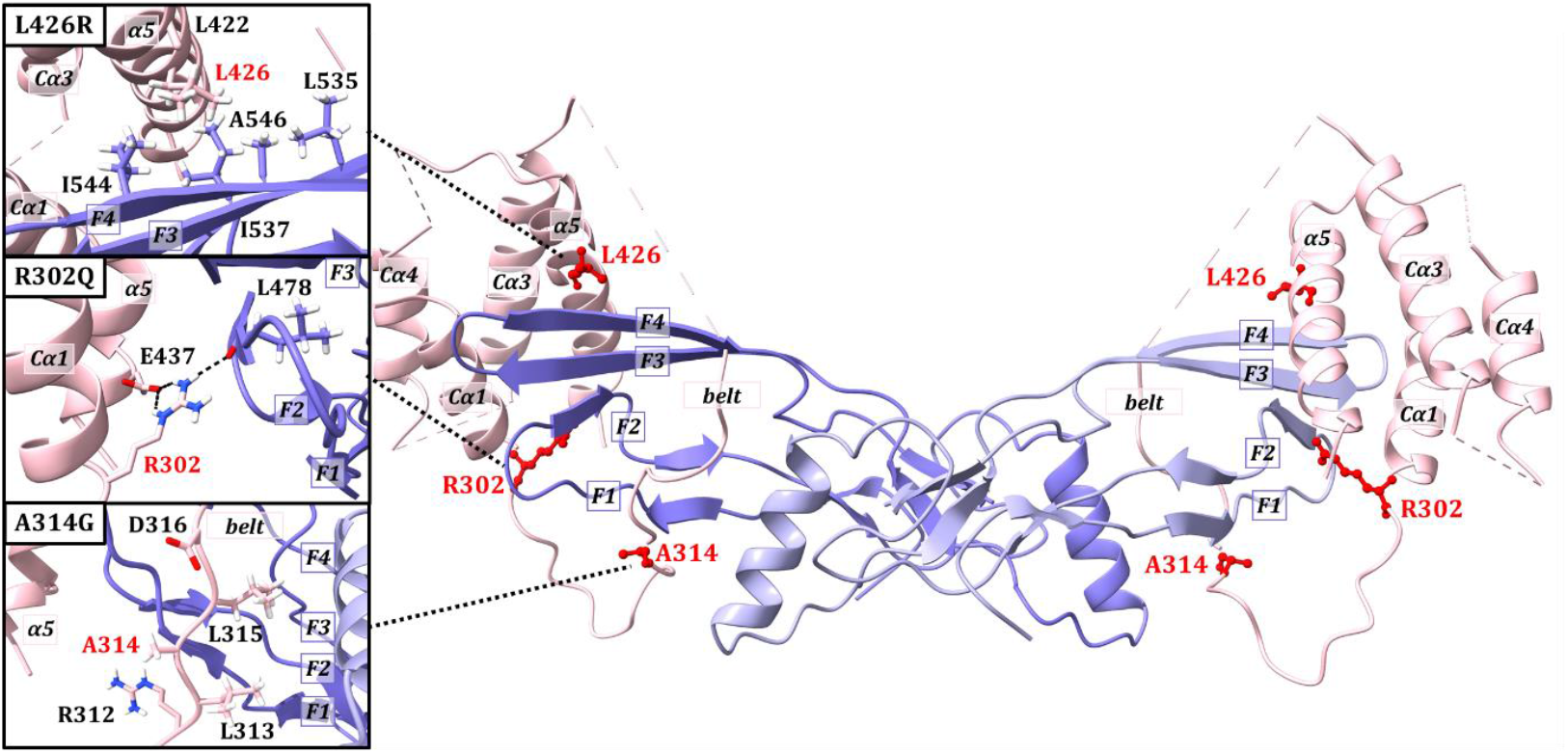
PMDS mutations within the HBD. AMH GF displayed with mutated amino acids in the HBD interface depicted with a red ball-and-stick representation. The inset figures on the left illustrate the detailed environment around mutated residues: (top) the hydrophobic pocket in which (α5)-L426 is located; (middle) (Cα1)-R302···(F2)-L478 hydrogen bond, and (Cα1)-R302··· (α5)-E437 salt bridge; (bottom) orientation and environment of the side chains at the bottom of the binding belt, including A314.

**Extended Data Fig. 9.**
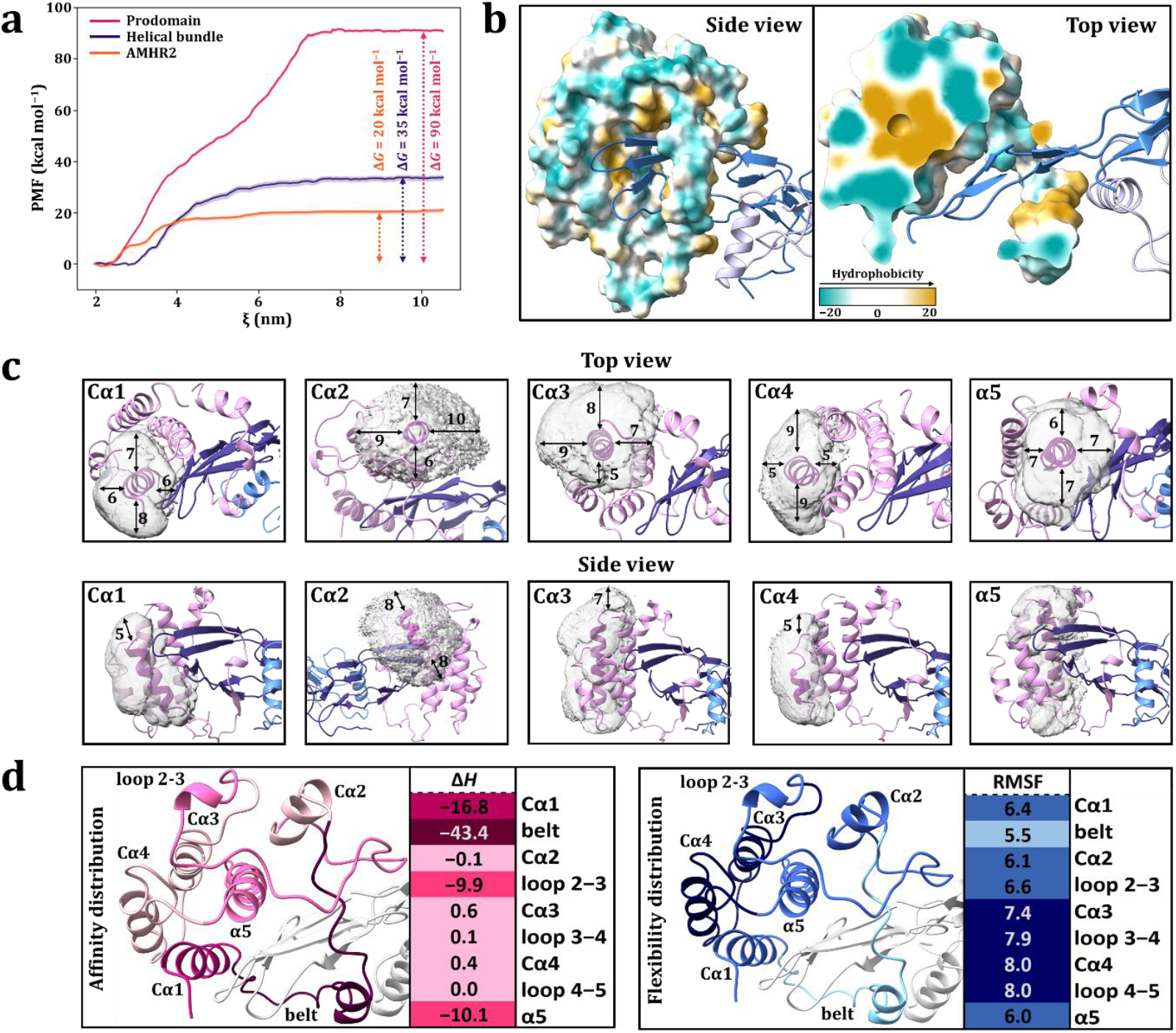
Computational analysis of binding energies and dynamics of the HBD:GF complex. **a**, Curves of the potential of mean force along reaction coordinate calculated from umbrella sampling simulations for the dissociation of the full prodomain HBD (in pink), HBD without the binding belt (in blue), and AMHR2 (in orange) from the GF with the corresponding Δ*G* values. The prodomain HBD profile is biphasic: helical bundle separates with an energy cost of 40 kcal mol^−1^, followed by the belt lagging behind, requiring an additional 50 kcal mol^−1^ for complete dissociation. This approximately uniform energy distribution has an unequal size distribution, as the binding belt constitutes only 17% of the amino acid sequence of the HBD. **b**, The left image shows the hydrophobic contacts with the helical bundle formed on the convex surface of the GF, while the right shows the top view of a cross-section through the hydrophobic surface that reveals the diffuse non-polar core of the HBD. **c**, 3D histograms of the positions of individual α-helices within the HBD during MD simulation. The range of axial and lateral motion in Å is shown together with the average structure as a reference. **d**, The decomposition of the binding energy on the left reveals an uneven distribution of the HBD-GF affinity: the binding belt (dark pink) constitutes 50% of the total prodomain contribution, followed by Cα1 (medium pink) at 20%, and loop 2-3 and α5 (pink) each at 12%. The remaining regions contribute insignificantly. According to the RMSF values, distribution of flexibility shows that Cα3 and Cα4, together with the corresponding loops represent the most dynamic parts (dark blue), while the belt shows the highest conformational stability (light blue) (**Supplementary Table 5, Supplementary Fig. 3**).

