## Supplemental Material for "Structural Basis of Non-Latent Signaling by the Anti-Müllerian Hormone Procomplex"

[1]Department of Pharmacology & Systems Physiology, University of Cincinnati, Cincinnati, OH, United States. [2]Department of Molecular & Cellular Biosciences, University of Cincinnati, Cincinnati, OH, United States. [3]Department of Chemistry, Boston University, Boston, MA, United States. [4]Pediatric Surgical Research Laboratories, Massachusetts General Hospital, Department of Surgery, Harvard Medical School, Boston, MA, United States.

### Table of Contents

| Content | Pages |
|---|---|
| Supplementary Tables 1−5 | S2–S6 |
| Supplementary Figs. 1−3 | S7–S9 |

**Supplementary Table 1 |** The MM-GBSA binding enthalpies for the AMH GF bound to HBD and AMHR2. The energies are divided into individual contributions of interacting components: helical bundles and binding belts in the case of HBD, and separate extracellular domains in the case of AMHR2. The simulations of both systems were carried out in five replicas and the calculated values are given in kcal mol$^{-1}$.

| | | **Binding Enthalpies (kcal mol$^{-1}$)** | | | | | | |
|---|---|---|---|---|---|---|---|---|
| | | **#1** | **#2** | **#3** | **#4** | **#5** | **AVG** | **STD** |
| | **Δ*H* of four semi–independent motifs of the prodomain HBD** | | | | | | | |
| 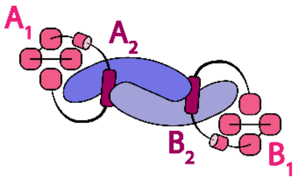 | $A_1$ | −25.28 | −34.71 | −57.73 | −49.41 | −39.32 | **−41.29** | *11.31* |
| | $A_2$ | −58.83 | −48.23 | −35.01 | −45.31 | −44.69 | **−46.41** | *7.64* |
| | $B_1$ | −39.31 | −37.62 | −35.73 | −42.30 | −48.92 | **−40.78** | *4.61* |
| | $B_2$ | −29.87 | −42.39 | −47.94 | −42.60 | −38.63 | **−40.29** | *5.99* |
| | AVG (**1**) | | | | | | **−41.04** | |
| | AVG (**2**) | | | | | | **−43.35** | |
| | **Δ*H* of one AMHR2 molecule** | | | | | | | |
| 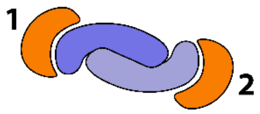 | ECD (**1**) | −33.10 | −40.83 | −35.52 | −33.29 | −45.75 | **−37.70** | *4.90* |
| | ECD (**2**) | −37.76 | −29.75 | −36.81 | −40.65 | −42.66 | **−37.53** | *4.41* |
| | AVG | | | | | | **−37.61** | |

**Supplementary Table 2 |** Number of hydrogen bonds between ligand fingers 2···3 and 3···4 forming β-sheets during MD simulations of the procomplex. The simulations were performed in five replicas, each 300 ns long, and the number of counted hydrogen bonds between marked pairs of amino acids is given for each monomer separately. The percentage (in red) is calculated based on full trajectories consisting of 150,000 frames.

**Number of hydrogen bonds in the procomplex**

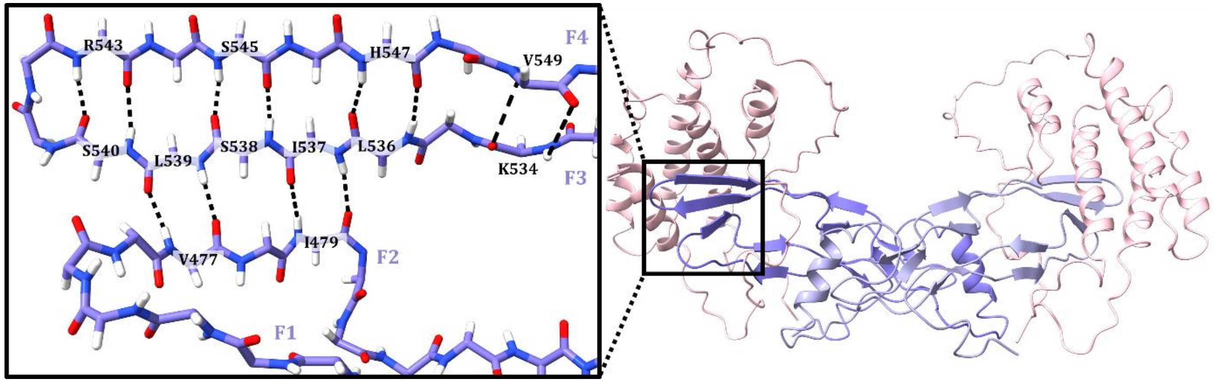


| | FINGERS 4···3 | | | | | | | | FINGERS 3···2 | | | |
|---|---|---|---|---|---|---|---|---|---|---|---|---|
| **AMH (A)** | **R543:N S540:O** | **R543:O S540:N** | **S545:N S538:O** | **S545:O S538:N** | **H547:N L536:O** | **H547:O L536:N** | **V549:N K534:O** | **V549:O K534:N** | **L539:O V477:N** | **L539:N V477:O** | **I537:O I479:N** | **I537:N I479:O** |
| **#1** | 8275 | 55641 | 94901 | 121044 | 110831 | 112719 | 47707 | 37053 | 117748 | 106159 | 108583 | 80271 |
| **#2** | 5257 | 45014 | 95388 | 113432 | 108562 | 132223 | 52393 | 23920 | 105712 | 106674 | 120173 | 32084 |
| **#3** | 4245 | 35469 | 107702 | 111480 | 66856 | 109794 | 36937 | 14826 | 107830 | 110364 | 83742 | 8111 |
| **#4** | 8985 | 70565 | 77439 | 111756 | 80873 | 110944 | 43348 | 19360 | 94914 | 106068 | 105023 | 36819 |
| **#5** | 7056 | 46383 | 88560 | 119273 | 78388 | 108206 | 32327 | 17078 | 107387 | 104543 | 95307 | 38009 |
| **AVG** | **6764** | **50614** | **92798** | **115397** | **89102** | **114777** | **42542** | **22447** | **106718** | **106762** | **102566** | **39059** |
| **%** | **5** | **34** | **62** | **77** | **59** | **77** | **28** | **15** | **71** | **71** | **68** | **26** |
| SUM | 534441 | | | | | | | | 355105 | | | |
| **AMH (B)** | **R543:N S540:O** | **R543:O S540:N** | **S545:N S538:O** | **S545:O S538:N** | **H547:N L536:O** | **H547:O L536:N** | **V549:N K534:O** | **V549:O K534:N** | **L539:O V477:N** | **L539:N V477:O** | **I537:O I479:N** | **I537:N I479:O** |
| **#1** | 8414 | 47943 | 92189 | 113968 | 93633 | 124038 | 32795 | 44843 | 107366 | 107422 | 91592 | 7895 |
| **#2** | 15182 | 61592 | 88015 | 129413 | 60881 | 106396 | 92084 | 27381 | 114586 | 95658 | 117694 | 51495 |
| **#3** | 6398 | 32551 | 84242 | 117253 | 60490 | 122838 | 58057 | 7935 | 80489 | 110721 | 80275 | 1136 |
| **#4** | 14768 | 48129 | 92654 | 124725 | 109054 | 94238 | 27069 | 31705 | 103072 | 100860 | 115616 | 49339 |
| **#5** | 15464 | 55691 | 58635 | 130830 | 100465 | 59921 | 6 | 7 | 126155 | 103082 | 112548 | 95373 |
| **AVG** | **12045** | **49181** | **83147** | **123238** | **84905** | **101486** | **42002** | **22374** | **106334** | **103549** | **103545** | **41048** |
| **%** | **8** | **33** | **55** | **82** | **57** | **68** | **28** | **15** | **71** | **69** | **69** | **27** |
| SUM | 518378 | | | | | | | | 354476 | | | |
| **AVG SUM** | **526410** | | | | | | | | **354791** | | | |

**Supplementary Table 3 |** Number of hydrogen bonds between ligand fingers 2···3 and 3···4 forming β-sheets during MD simulations of the AMH-AMHR2 complex. The simulations were performed in five replicas, each 300 ns long, and the number of counted hydrogen bonds between marked pairs of amino acids is given for each monomer separately. The percentage (in red) is calculated based on full trajectories consisting of 150,000 frames.

**Number of hydrogen bonds in the AMH-AMHR2 complex**

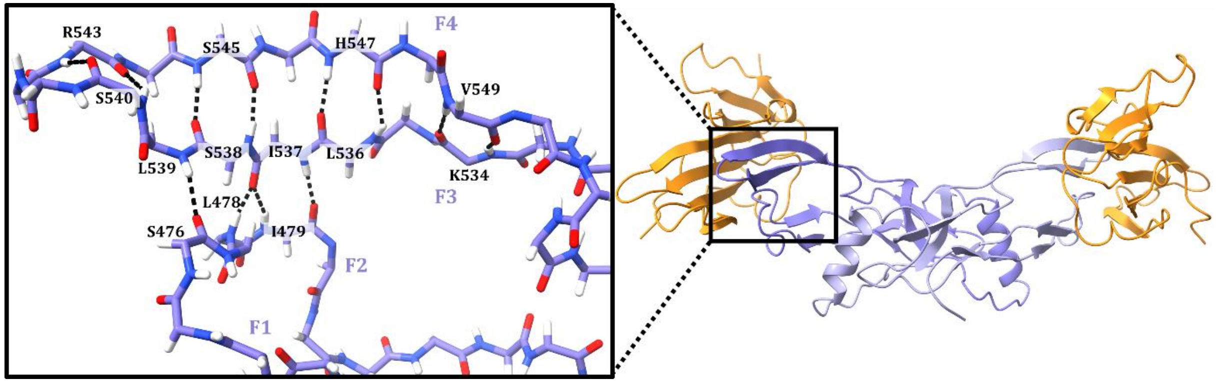


| | FINGERS 4···3 | | | | | | | | FINGERS 3···2 | | | |
|---|---|---|---|---|---|---|---|---|---|---|---|---|
| **AMH (A)** | **R543:N S540:O** | **R543:O S540:N** | **S545:N S538:O** | **S545:O S538:N** | **H547:N L536:O** | **H547:O L536:N** | **V549:N K534:O** | **V549:O K534:N** | **Leu539:N Ser476:O** | **Ile537:O Leu478:N** | **Ile537:O Ile479:N** | **Ile537:N Ile479:O** |
| **#1** | 4695 | 41774 | 90868 | 107711 | 59152 | 104483 | 73021 | 82211 | 97416 | 132557 | 6992 | 126752 |
| **#2** | 18803 | 104409 | 125568 | 90442 | 50300 | 79580 | 65813 | 80659 | 91654 | 125135 | 12626 | 124463 |
| **#3** | 7683 | 66694 | 123100 | 102129 | 50020 | 85099 | 71565 | 84004 | 91834 | 128125 | 9718 | 128255 |
| **#4** | 569 | 69984 | 126757 | 98477 | 48057 | 80935 | 69297 | 85881 | 84503 | 128124 | 8202 | 125413 |
| **#5** | 8770 | 81634 | 128782 | 92444 | 56697 | 79429 | 62835 | 82089 | 85791 | 133704 | 8039 | 119759 |
| **AVG** | **8104** | **72899** | **119015** | **98241** | **52845** | **85905** | **68506** | **82969** | **90240** | **129529** | **9115** | **124928** |
| **%** | **5** | **49** | **79** | **65** | **35** | **57** | **46** | **55** | **60** | **86** | **6** | **83** |
| SUM | 588484 | | | | | | | | 353812 | | | |
| **AMH (B)** | **R543:N S540:O** | **R543:O S540:N** | **S545:N S538:O** | **S545:O S538:N** | **H547:N L536:O** | **H547:O L536:N** | **V549:N K534:O** | **V549:O K534:N** | **Leu539:N Ser476:O** | **Ile537:O Leu478:N** | **Ile537:O Ile479:N** | **Ile537:N Ile479:O** |
| **#1** | 2463 | 35490 | 125959 | 100173 | 36982 | 65378 | 70825 | 79808 | 102923 | 129055 | 8806 | 128466 |
| **#2** | 3435 | 13344 | 114031 | 95796 | 58480 | 89596 | 74329 | 91136 | 88205 | 129480 | 9548 | 115548 |
| **#3** | 26752 | 107018 | 120734 | 99260 | 65656 | 97647 | 86329 | 96135 | 103139 | 132492 | 16038 | 110732 |
| **#4** | 22193 | 107380 | 122077 | 90264 | 61857 | 85702 | 67616 | 85079 | 93194 | 131351 | 10992 | 117914 |
| **#5** | 5951 | 59806 | 109837 | 97942 | 46343 | 73928 | 72370 | 85013 | 92525 | 123131 | 10271 | 123279 |
| **AVG** | **12159** | **64608** | **118528** | **96687** | **53864** | **82450** | **74294** | **87434** | **95997** | **129102** | **11131** | **119188** |
| **%** | **8** | **43** | **79** | **64** | **36** | **55** | **50** | **58** | **64** | **86** | **7** | **79** |
| SUM | 590024 | | | | | | | | 355418 | | | |
| **AVG SUM** | **589254** | | | | | | | | **354615** | | | |

**Supplementary Table 4 |** Average RMSD values of the ligand backbone atoms during 300 ns-MD simulations of the procomplex (prodomain-HBD) and the AMH-AMHR2(ECD) complex. The simulations were carried out in five replicas, and the calculated values are given Å.

| | | | | | | |
|---|---|---|---|---|---|---|
| | | | **Procomplex** | | | |
| **#1** | **#2** | **#3** | **#4** | **#5** | **AVG** | **STD** |
| 3.59 | 3.72 | 4.41 | 3.88 | 3.97 | **3.91** | *0.28* |
| | | | **AMH-AMHR2** | | | |
| **#1** | **#2** | **#3** | **#4** | **#5** | **AVG** | **STD** |
| 1.75 | 1.75 | 1.89 | 1.91 | 1.74 | **1.81** | *0.08* |

**Supplementary Table 5 |** The decomposition of the binding enthalpy into individual components of the prodomain was calculated on 3,000 frames during the last 30 ns of each 300 ns-long MD simulation of the procomplex. The average per-residue RMSF values of individual components of the prodomain were calculated on the last 30 ns of the trajectories in order to make a comparison with the calculated energy contributions. The plotted average RMSF values are shown in the **Supplementary Fig. 3**. The simulations were performed in five replicas, and the calculated values are shown for each monomer separately. In the last column, the average values of enthalpy and RMSF for both monomers are given, and the intensity of the color corresponds to the affinity or flexibility.

| | | **Enthalpy (kcal mol⁻¹)** | | | | | | | **RMSF (Å)** | | | | | | |
|---|---|---|---|---|---|---|---|---|---|---|---|---|---|---|---|
| | **Resid** | **#1** | **#2** | **#3** | **#4** | **#5** | **AVG** | **STD** | **#1** | **#2** | **#3** | **#4** | **#5** | **AVG** | **STD** |
| **Prodom. (A)** | **287–451** | −84.11 | −82.94 | −92.74 | −94.72 | −84.01 | **−87.70** | *4.98* | 6.67 | 8.66 | 5.66 | 4.97 | 7.13 | **6.62** | *1.27* |
| **Cα1** | **287–302** | −17.15 | −18.58 | −11.33 | −13.88 | −18.39 | **−15.87** | *2.83* | 5.54 | 7.19 | 5.48 | 4.48 | 7.57 | **6.05** | *1.15* |
| **belt** | **303–331** | −58.83 | −48.23 | −35.01 | −45.31 | −44.69 | **−46.41** | *7.64* | 4.32 | 6.71 | 4.82 | 4.77 | 6.68 | **5.46** | *1.03* |
| **Cα2** | **332–342** | −0.08 | 0.33 | 0.10 | 0.08 | 0.30 | **0.15** | *0.15* | 6.77 | 8.47 | 4.48 | 4.68 | 4.93 | **5.87** | *1.54* |
| **loop 2–3** | **343–371** | −3.14 | −8.71 | −15.62 | −18.97 | −6.22 | **−10.53** | *5.89* | 8.13 | 8.97 | 5.78 | 4.76 | 4.93 | **6.51** | *1.72* |
| **Cα3** | **372–393** | 0.50 | 0.51 | 0.64 | 0.55 | 0.59 | **0.56** | *0.05* | 7.93 | 10.41 | 6.16 | 5.48 | 7.81 | **7.56** | *1.71* |
| **loop 3–4** | **394–399** | 0.10 | 0.09 | 0.06 | 0.10 | 0.08 | **0.09** | *0.01* | 6.33 | 10.18 | 7.12 | 5.89 | 10.94 | **8.09** | *2.07* |
| **Cα4** | **400–411** | 0.53 | 0.36 | 0.39 | 0.36 | 0.34 | **0.40** | *0.07* | 7.69 | 9.86 | 7.16 | 5.91 | 10.09 | **8.14** | *1.61* |
| **loop 4–5** | **412–419** | 0.06 | 0.13 | 0.09 | 0.16 | −1.19 | **−0.15** | *0.52* | 9.56 | 10.60 | 7.63 | 5.73 | 7.56 | **8.21** | *1.70* |
| **α5** | **420–441** | −7.61 | −9.82 | −17.67 | −10.73 | −9.78 | **−11.12** | *3.43* | 6.35 | 8.18 | 5.30 | 4.26 | 6.38 | **6.09** | *1.30* |
| **Prodom. (B)** | **287–451** | −69.18 | −80.01 | −83.67 | −84.90 | −87.55 | **−81.06** | *6.42* | 5.99 | 8.50 | 5.93 | 5.14 | 7.42 | **6.60** | *1.20* |
| **Cα1** | **287–302** | −19.01 | −24.27 | −17.75 | −15.72 | −12.39 | **−17.83** | *3.92* | 6.25 | 8.15 | 6.60 | 5.32 | 7.83 | **6.83** | *1.04* |
| **belt** | **303–331** | −29.87 | −42.39 | −47.94 | −42.60 | −38.63 | **−40.29** | *5.99* | 5.91 | 7.32 | 4.91 | 4.20 | 5.50 | **5.57** | *1.05* |
| **Cα2** | **332–342** | −0.16 | 0.16 | −0.54 | 0.08 | −1.60 | **−0.41** | *0.64* | 4.93 | 8.70 | 4.90 | 5.70 | 7.80 | **6.41** | *1.56* |
| **loop 2–3** | **343–371** | −12.1 | −7.74 | −9.05 | −2.16 | −15.07 | **−9.22** | *4.35* | 4.95 | 8.75 | 5.44 | 6.17 | 7.85 | **6.63** | *1.44* |
| **Cα3** | **372–393** | 0.65 | 0.72 | 0.50 | 0.58 | 0.59 | **0.61** | *0.07* | 6.43 | 9.53 | 6.58 | 5.37 | 8.47 | **7.27** | *1.51* |
| **loop 3–4** | **394–399** | 0.07 | 0.09 | 0.07 | 0.07 | 0.07 | **0.07** | *0.01* | 7.89 | 11.13 | 7.33 | 4.86 | 7.68 | **7.78** | *2.00* |
| **Cα4** | **400–411** | 0.35 | 0.52 | 0.38 | 0.29 | 0.32 | **0.37** | *0.08* | 7.27 | 9.94 | 7.63 | 5.33 | 8.67 | **7.77** | *1.53* |
| **loop 4–5** | **412–419** | 0.16 | 0.05 | 0.18 | 0.11 | 0.13 | **0.13** | *0.04* | 6.19 | 8.86 | 7.57 | 6.27 | 9.90 | **7.76** | *1.45* |
| **α5** | **420–441** | −7.44 | −6.61 | −6.79 | −11.50 | −12.91 | **−9.05** | *2.63* | 5.43 | 7.05 | 5.62 | 4.45 | 7.15 | **5.94** | *1.03* |

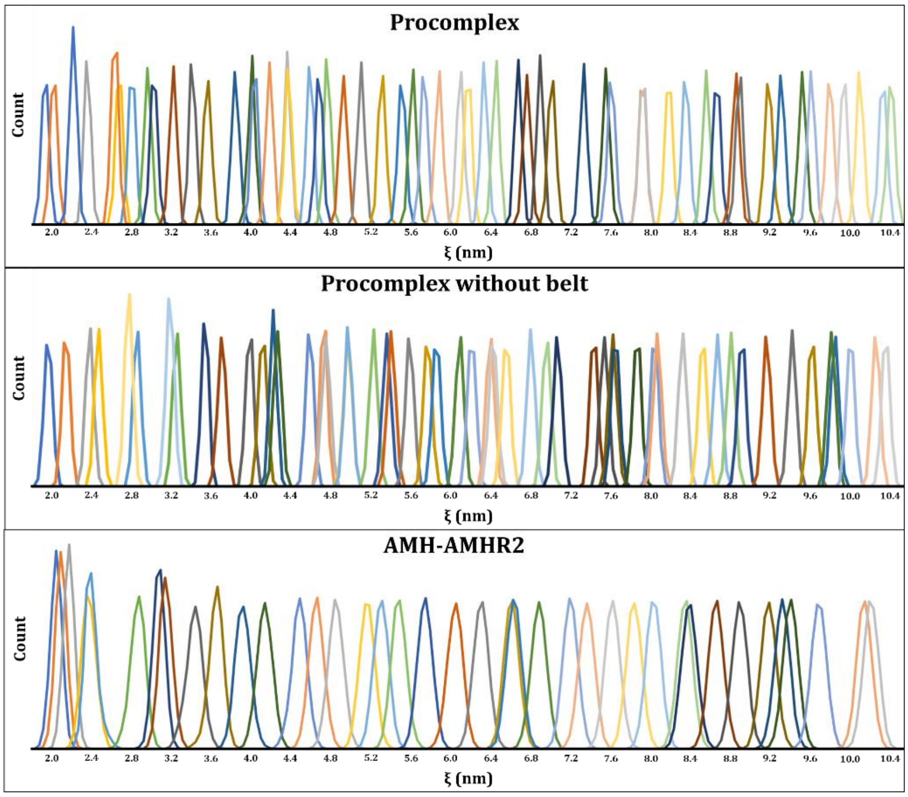


**Supplementary Fig. 1 |** Histograms depicting the procomplex (top), procomplex without the belt (middle), and AMH-AMHR2 (bottom) show the probability distribution of the distance between the centers of mass of AMH and the prodomain or receptor along the reaction coordinate for each window of umbrella sampling simulations.

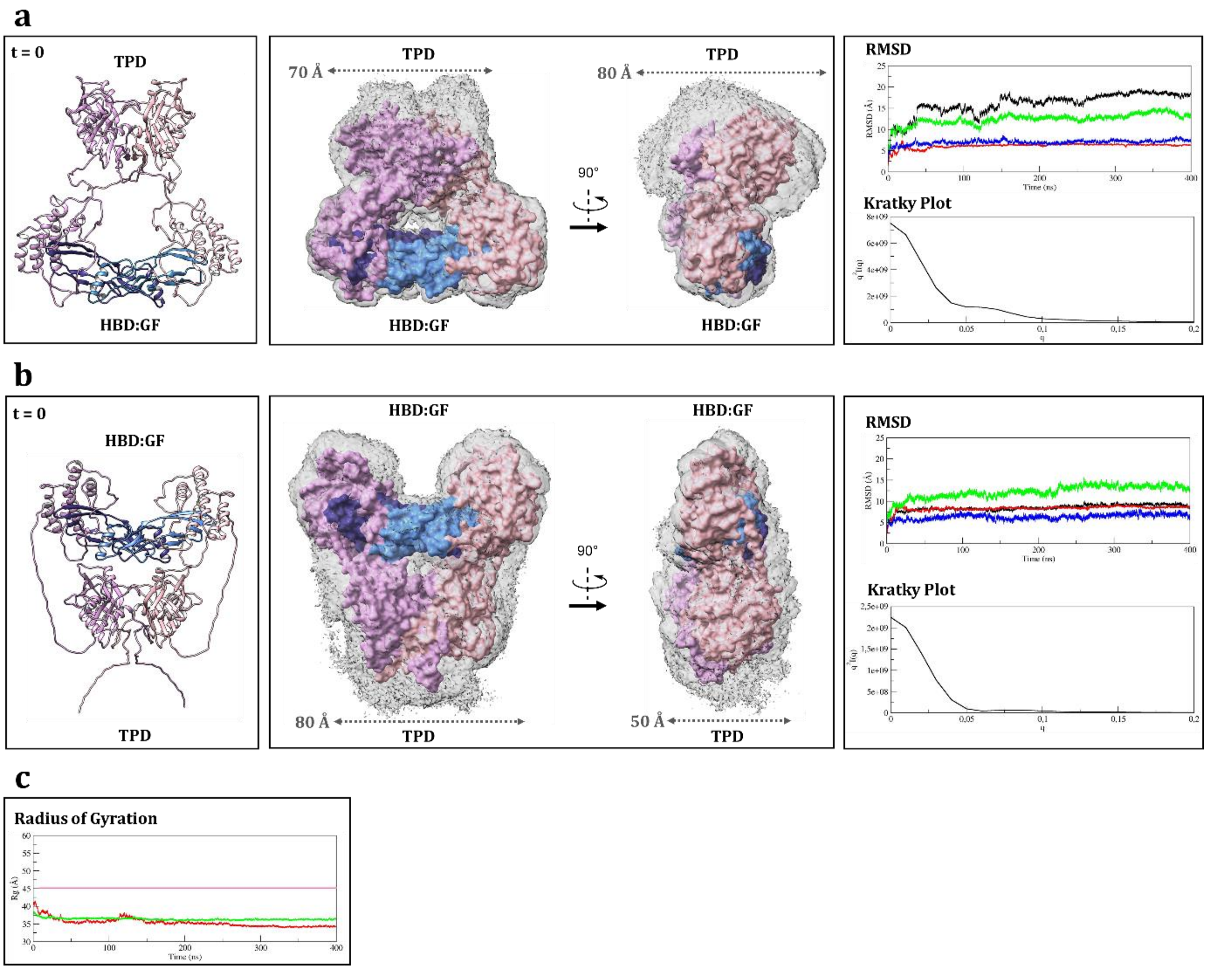


**Supplementary Fig. 2 |** Two additional AlphaFold2-based starting models for MD simulation. **a**, model02; **b**, model03. On the Left is a representation of their initial structures at time t = 0, 3D histograms of the procomplex position during MD simulations shown together with the average structure as a reference in the middle, and RMSD graphs as well as Kratky plots from SAXS_MD analysis are shown on the right. Prodomains are colored pink and purple, and monomers in the GF are represented in different shades of blue. On the RMSD graphs, the black line refers to the backbone atoms of the entire procomplex, while red, green and blue lines pertain to individual parts of the prodomain: TPD, linkers and HBD, respectively. **c**, Evolution of the radius of gyration during MD simulations for model02 (in red), and model03 (in green), together with the experimental value of 45.5 Å shown as a pink line.

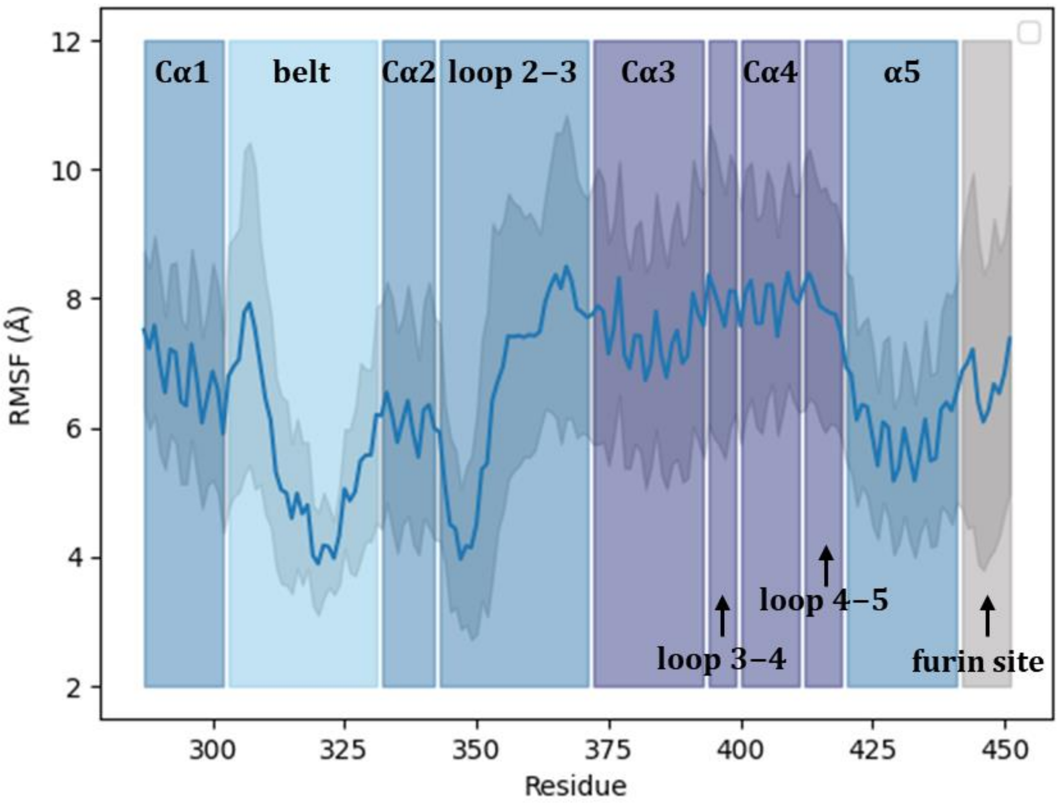


**Supplementary Fig. 3 |** Average RMSF values of both monomers of the prodomain calculated on the last 30 ns of trajectories through five replicas. The individual components of the prodomain are indicated on the graph, and the color intensity corresponds to the degree of flexibility. The shaded region surrounding the average value represents the standard deviation across five replicates.
